# Multicentre comparison of biological and functional properties of mesenchymal stromal cells from different sources cultivated using a harmonised manufacturing workflow

**DOI:** 10.1101/2022.09.14.507944

**Authors:** Sandra Calcat-i-Cervera, Erika Rendra, Eleonora Scaccia, Francesco Amadeo, Vivien Hanson, Bettina Wilm, Patricia Murray, Timothy O’Brien, Arthur Taylor, Karen Bieback

**Author notes:** **Corresponding author** Prof. Dr. rer. nat. Karen Bieback, Institute of Transfusion Medicine and Immunology, Medical Faculty Mannheim, Heidelberg University; German Red Cross Blood Service Baden-Württemberg – Hessen, Friedrich-Ebert Str. 107, D-68167 Mannheim. Authors contributed equally (Sandra Calcat-i-Cervera, Erika Rendra, Eleonora Scaccia, Francesco Amadeo, Arthur Taylor, Karen Bieback).

## Abstract

**Background:** Mesenchymal stromal cells (MSCs), commonly sourced from adipose tissue, bone marrow and umbilical cord, have been widely used in many medical conditions due to their therapeutic potential. Yet, the still limited understanding of the underlying mechanisms of action hampers clinical translation. Clinical potency can vary considerably depending on tissue source, donor attributes, but importantly, also culture conditions. Lack of standard procedures hinders inter-study comparability and delays the progression of the field. The aim of this study was A-to assess the impact on MSC characteristics when different laboratories performed analysis on the same MSC material using harmonised culture conditions and B-to understand source-specific differences.

**Methods:** Three independent institutions performed a head-to-head comparison of human-derived adipose (A-), bone marrow (BM-), and umbilical cord (UC-) MSCs using harmonised culture conditions. In each centre, cells from one specific tissue source were isolated and later distributed across the network to assess their biological properties, including cell expansion, immune phenotype, and tri-lineage differentiation (part A). To assess tissue specific function, angiogenic and immunomodulatory properties and the *in vivo* biodistribution were compared in one expert lab (part B).

**Results:** By implementing a harmonised manufacturing workflow, we obtained largely reproducible results across three independent laboratories in part A of our study. Unique growth patterns and differentiation potential were observed for each tissue source, with similar trends observed between centres. Immune phenotyping verified expression of typical MSC surface markers and absence of contaminating surface markers. Depending on the established protocols in the different laboratories, quantitative data varied slightly. Functional experiments in part B concluded that conditioned media from BM-MSCs significantly enhanced tubulogenesis and endothelial migration *in vitro*. In contrast, immunomodulatory studies reported superior immunosuppressive abilities for A-MSCs. Biodistribution studies in healthy mice showed lung entrapment after administration of all three types of MSCs, with a significantly faster clearance of BM-MSCs.

**Conclusion:** These results show the heterogeneous behaviour and regenerative properties of MSCs as a reflection of intrinsic tissue-origin properties while providing evidence that the use of standardised culture procedures can reduce but not eliminate inter-lab and operator differences.

**Highlights:** In this study, we have:

- Provided a harmonised manufacturing workflow that has demonstrated reproducible results across three independent laboratories when expanding MSCs.
- Defined a multi-assay matrix capable of identifying functional differences in terms of angiogenesis, wound healing abilities and immunosuppressive properties.
- Demonstrated similar *in vivo* biodistribution properties regardless of cell origin.

## Introduction

Mesenchymal stromal cells (MSCs) are multipotent cells that have attracted huge interest in different areas of regenerative medicine. Because of their unique immunomodulatory, anti-inflammatory and pro-regenerative abilities [1-3], their ease of isolation from multiple tissues [4] and high expansion potential ex vivo, MSCs have been extensively studied in several pre-clinical models and early-phase clinical trials to treat a variety of human diseases [1, 5].

MSCs were first isolated from bone marrow (BM-) in 1968 by Friedenstein *et al*. [6] and since then cells with similar properties have been identified in several other tissues (e.g. adipose tissue, umbilical cord, skin tissue) [4]. Although BM-MSCs are the most commonly used cell source in clinical trials [7], adipose (A-) and umbilical cord (UC-) derived MSCs have become quite attractive sources as they can be easily obtained with relatively good yields and less invasively [8]. The possibility to isolate MSCs from different starting materials elicits the question of whether it is more advantageous to use autologous or allogeneic cells. The use of autologous MSCs guarantees an easy source that does not evoke allo-immunity. However, it is associated with high costs of isolation, expansion, safety testing and donor-related comorbidities that might impact product quality [7]. Allogeneic cells may offer a more cost-effective and better standardisable off-the-shelf product. Hence, merging knowledge about basic cell characteristics (viability, proliferation, immunophenotype) together with bioactivity in a range of assays could help in identifying the ‘right’ source for the ‘right’ application.

Unfortunately, despite years of research and highly promising preclinical data, the translation to the clinic is well below expectations. In many clinical trials, MSCs have shown little benefit [1, 5, 9]. Inconsistent and poorly defined manufacturing procedures increase the heterogeneity that intrinsically exists in a field where donor variability and tissue origin have a strong role. Thus, when considering clinical translation, defining an optimal scalable manufacturing workflow is key to ensure product quality while minimising costs and timelines [10]. Numerous different manufacturing workflows have been established, which largely affect cell characteristics. Stroncek and colleagues recently demonstrated that variations in cell culture procedures affected the functional and molecular characteristics of the cells to a much higher extent than the source material itself, which was shipped across five different manufacturing centres [11]. The variation in culture conditions included the use of different media (type and composition), sera (origin and concentration in the medium) and seeding densities [11]. This emphasised that clinical-scale manufacturing requires optimisation, and importantly, worldwide standardisation. Within the context of the RenalToolBox EU ITN Network [https://www.renaltoolbox.org], which includes several leading EU academic institutions and industry experts, researchers from the University of Liverpool (Liverpool, UK), the University of Heidelberg (Heidelberg, Germany) and the University of Galway (Galway, Ireland) collaborated to assess biological and therapeutic properties of MSCs derived from bone marrow (BM-MSC), adipose tissue (A-MSC), and umbilical cord (UC-MSC) in a multi-centre comparative study. In part A of our study, we focused on comparing cell characteristics across centres using harmonised cultures conditions for A-, BM- and UC-MSCs mimicking three decentralised manufacturing sites. MSCs were generated in one centre, shipped as cryo-aliquots to the other centres and cultivated under harmonised standard culture conditions to compare cell behaviour, differentiation potential and expression of MSC markers *in vitro* (Figure 1).

**Figure 1.**
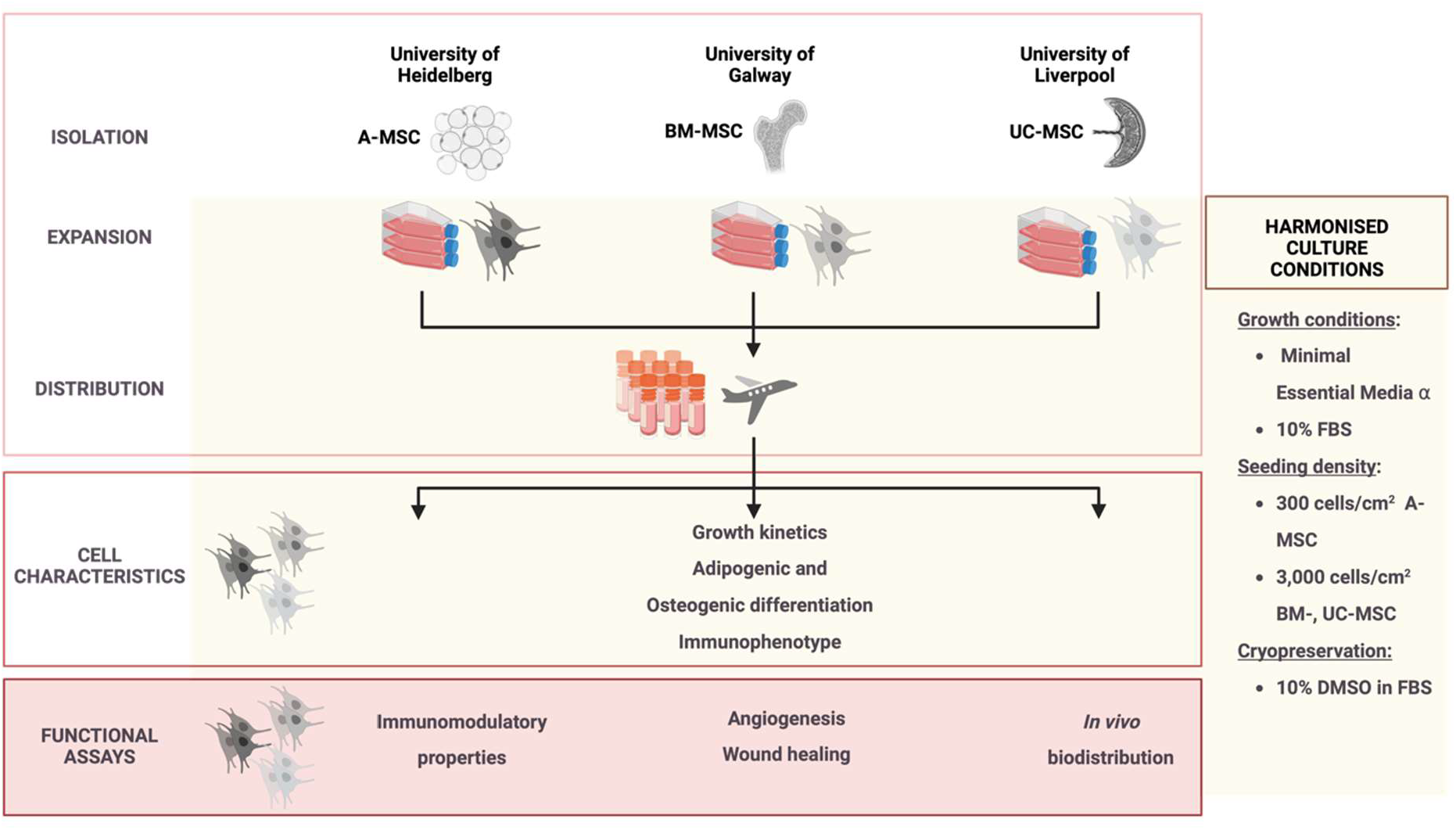
Schematic representing study design and assay distribution across centres.

Assessing the impact of harmonised manufacturing methods on biological properties beyond basic cell characterisation could provide helpful insights to decipher particular mechanisms of action of different tissue-origin MSCs. Thus, in part B, we assessed tissue source specificities further. Given that some of their therapeutic properties are elicited by their ability to release soluble bioactive factors to promote angiogenesis as well as to modulate immune responses [12-14], these properties were assessed individually in Galway and Heidelberg, respectively. The team in Liverpool compared the *in vivo* biodistribution in a small rodent model (Figure 1).

## Materials and Methods

### Mesenchymal Stromal Cells Culture

MSCs were obtained from different sites participating in the RenalToolBox network. A-MSCs from lipoaspirates were processed in Heidelberg after obtaining informed consent (Mannheim Ethics Commission; vote number 2006-192NMA). BM-MSCs provided by Galway were purchased from Lonza (Basel, Switzerland), and UC-MSCs with informed consent obtained in accordance with the Declaration of Helsinki were sourced from the NHS Blood and Transplant and transferred to the University of Liverpool. Three different donors per tissue source were isolated in each centre according to their standard procedures ([15] Galway, [16] Heidelberg). From passage 3 (A- and BM-MSCs) or passage 4 (UC-MSCS) on, cells were expanded using harmonised conditions (see supplementary data), and banked prior to distribution across the network (see below). After shipment and subsequent storage in liquid nitrogen, MSCs were thawed and cultivated under defined harmonised conditions. These included the basic growth medium (MEM-α media, Gibco, ThermoFisher Scientific, 2561029), a common lot of foetal bovine serum (FBS, Gibco, ThermoFisher Scientifics, 10270-106, Lot 42Q7096K) and optimised seeding densities (300 cells/cm^2^ for A-MSCs and 3,000 cells/cm^2^ for BM- and UC-MSCs) at 37 °C with 5% (v/v) CO_2_ and controlled humidity (see **supplementary data** for more information on FBS batch testing and seeding densities). All experiments were performed within a similar passage number, ranging from p4 to p6 depending on experimental requirements and intrinsic factors such as initial availability.

### Cryopreservation

Upon reaching 70% confluency, MSCs were cryopreserved for distribution across sites. Upon cell dissociation by trypsinisation (0.25%Trypsin-Ethylenediaminetetraacetic Acid 1X, Gibco, ThermoFisher Scientific, 25200-056), the cells were counted using appropriate methods (NucleoCounter NC-200 automated cell counter (Galway), CASY cell counter with dead cell exclusion (Heidelberg), manual cell counting (Liverpool)) and centrifuged. 5 × 10^5^ to 1 × 10^6^ cells/ml were resuspended in freezing media (FBS + 10% Dimethyl Sulfoxide, DMSO, Sigma, D2660) and frozen down.

### Conditioned media collection

Conditioned media (CM) were generated from MSCs at passage 4 to 6. Upon reaching 80% confluency, cells were washed with 1X DPBS and incubated for 24 hours in serum-free MEM-α media. The supernatant was collected and centrifuged for 5 minutes at 400 g to remove cell debris before being stored at –80 °C until further use.

### Part A-Basic MSC Characterisation Growth kinetics

To study the growth kinetics of MSCs, the population doublings (PDs) and population doubling time (PDTs) were calculated by seeding 300 or 3,000 cells/cm^2^ (A-MSCs, and BM- and UC-MSCs, respectively) at the start of a passage and counting the number of cells harvested at the end of said passage after reaching 70% confluency. PDTs were calculated as PDT = t x log2/(log N_t_ – log N_0_) while PDs were calculated as PD = log2 (N_t_ / N_0_); t indicates time in culture, N_t_ the number of harvested cells and N_0_ the number of seeded cells.

### Adipogenic and osteogenic differentiation

Adipogenic and osteogenic potential of MSCs was obtained using commercially available media: Adipogenic Differentiation Medium 2 (PromoCell, C-28016) and Osteogenic Differentiation Medium (PromoCell, C-28013), respectively. Harvested MSCs were seeded at a density of 5,700 cells/well for adipogenesis and 2,900 cells/well for osteogenesis in cell culture treated 96-well plates and kept at 37 °C. After 48 hours, differentiation was induced by adding differentiation media to positive differentiated cultures while undifferentiated cells were kept in standard growth medium. Medium was replenished twice a week and differentiation assessed after 14 days.

Quantitative analysis of adipogenic and osteogenic differentiation was assessed using the AdipoRed™ Analysis Reagent (Lonza, PT-7009) and OsteoImage™ Mineralization (Lonza, PA-1503), respectively, as per the manufacturer’s instructions. For normalisation, cells were also stained with Hoechst 33342 (Invitrogen, 917368). The emitted fluorescent signal from adipogenic and osteogenic quantification and Hoechst staining were measured using a multimode plate reader. Data were presented as a fold-change of the undifferentiated cultures.

### Immunophenotypic analysis

Flow cytometry characterisation was performed in each centre according to their routinely used procedures and equipment (**Supplementary table 1**). MSCs were harvested when cell confluence was reached and resuspended in FACS buffer. Cells were stained at 4°C for 20 minutes and data was acquired using conventional flow cytometers. A minimum of 10^4^ events was analysed for each marker.

### Part B – Functional MSC Characterisation Angiogenic assays

#### Endothelial cell tube formation assay

Human umbilical cord endothelial vein cells (HUVECs, Lonza, C2519A) were grown in endothelial growth medium (EGM-2, Lonza, CC-3162) until 90% confluent. Further, 48-well plates were coated with 110 µl of growth-factor reduced Matrigel (Corning, 734-1101) and left to gel. HUVECs were harvested and resuspended in MSC-CM at a concentration of 25,000 cells/well. HUVECs stimulated with standard EGM-2 containing 10 ng/ml vascular endothelial growth factor (VEGF) served as positive controls, and cultures with MSC growth medium as negative controls. Plates were then incubated for 18 hours and all conditions were assessed in triplicates. A total of six images were acquired per well with a 4X lens on an Olympus CKX41 brightfield microscope fitted with HD Chrome camera (1/.8”) and 10x C-mount adapter and analysed using the angiogenesis analyser plugin for ImageJ (National Institutes of Health, Bethesda, USA).

#### Wound scratch assay

HUVECs were seeded in 48-well plates at 84,000 cells/cm^2^ and cultured overnight. Subsequently, a p200 tip was used to create a scratch in each monolayer. Cultures were washed with DPBS before adding MSC-CM. Scratches were imaged immediately after the addition of CM (0 hours) and after 8 and 24 hours incubation using the automated Cytation 1 Imaging Reader at 4X (BioTek, with Gen5 Version 3.04 software, Swindon, UK). Six replicates were undertaken, and the total area of each scratch was measured using Image J and the percentage of closure was calculated relative to time 0 hours.

#### Angiogenesis Cytokine Array

The relative levels of angiogenesis-related cytokines in the MSC-CM were analysed using the Proteome Profiler Human Cytokine Array Kit from R&D systems (Abingdon, UK, ARY022B) per manufacturer’s instructions. Levels of angiogenic cytokines are expressed relative to the internal control of each sample.

### Immunomodulatory assays

#### PBMC Proliferation Assay

MSC-mediated inhibition of T cell proliferation was assessed as described before [17]. MSCs were seeded one day before adding peripheral blood mononuclear cells (PBMCs) isolated from leukapheresis samples from healthy donors, provided by the German Red Cross Blood Donor Service in Mannheim (Mannheim Ethics Commission; vote number 2018-594N-MA). To assess their proliferation, PBMCs were labelled with proliferation dye Cytotell Green (ATT Bioquest, 22253) (1:500 dilution) and seeded at a 1:10 MSCs:PBMCs ratio in RPMI, supplemented with 10% FBS, 2% L-glutamine (PAN Biotech, P04-80100), 1% Penicillin/Streptomycin (PAN Biotech, P06-07100), and 200 U/ml IL-2 (Promokine, C61240). PBMC proliferation was stimulated with phytohemagglutinin-L (PHA, 4.8 µg/ml (Biochrom, Merck Millipore, M5030)). PBMCs cultured alone without MSCs in the absence and presence of PHA served as negative and positive controls, respectively.

After 5 days, PBMC proliferation was measured based on the dilution of Cytotell Green dye using a FACS Canto II (BD Biosciences) and the data were analysed with FlowJo Software.

#### IFN-γ stimulation and intracellular Indoleamine 2,3-dioxygenase (IDO) Staining

Indoleamine 2,3-dioxygenase (IDO)-mediated tryptophan degradation supresses T cell proliferation as described before [17]. To assess the level of IDO expression in MSCs, the cells were treated in the presence or absence of interferon γ (IFN-γ 25ng/ml (R&D Systems, 285-IF) for 24 hours. For intracellular IDO staining, MSCs were harvested, fixed, permeabilised and then stained (anti-IDO PE antibody (1:40 dilution) (ThermoFisher Scientific, 12-9477-42)). After washing, the cells’ fluorescence was measured with a FACS Canto (BD Biosciences) and the data analysed with FlowJo.

#### Biodistribution *in vivo*

Biodistribution of the different MSCs in mice was evaluated by bioluminescence imaging (BLI). For this purpose, the cells were transduced to express a firefly luciferase genetic reporter.

#### Production of FLuc^+^ expressing cells

MSCs were transduced with a lentiviral vector (LV) encoding the luc2 firefly luciferase (FLuc) reporter. The pHIV-Luc2-ZsGreen vector was a gift from Bryan Welm and Zena Werb (Addgene plasmid #39,196). The LV also contain a gene encoding for a green fluorescent protein, ZsGreen. Lentiviral particles were produced using standard protocols [18] by co-transfection of HEK cells with the transfer vector (pHIV-Luc2-ZsGreen or pHIV-AkaLuc-ZsGreen), an envelope plasmid (pMD2.G) and a packaging plasmid (psPAX2), concentration by ultracentrifugation and titration using HEK cells, based on ZsGreen expression.

To produce the transduced populations, MSCs were infected overnight with a multiplicity of infection of 5 in the presence of 6 μg/mL diethylaminoethyl-dextran (DEAE-dextran) [19]. The cells were then grown until 60-90% confluence before sorting based on ZsGreen fluorescence using a FACSaria II (BD Biosciences) to obtain a pure population of cells expressing the transgene (FLuc^+^ MSCs).

#### Animal experiments

7-9-week-old C57 Black 6 (C57BL/6) albino female mice were used to evaluate the biodistribution of FLuc^+^ MSCs from their administration into the animal (day 0) up to 7 days later. Mice were obtained from a colony managed by the Biomedical Services Unit at the University of Liverpool (UK). Mice were housed in individually ventilated cages under a 12-hour light/dark cycle and provided with standard food and water ad libitum. All animal procedures were performed under a licence granted under the UK’s Animals (Scientific Procedures) Act 1986 and were approved by the University of Liverpool Animal Welfare and Ethics Research Board.

FLuc^+^ MSCs were harvested and suspended in ice-cold DPBS at a concentration of 2.5×10^5^ cells/100 μL and kept on ice until administration. Animals (n = 4 per donor per cell type) were anaesthetised with isoflurane and intravenously (IV) injected with 100 μL of cell suspension through the tail vein, followed by subcutaneous administration (SC) of 200 μL of 47 mM D-Luciferin 20 minutes before imaging [20]. The administration of the substrate and the imaging were performed on the day of the injection of the cells (day 0) and after 1, 3 and 7 days. Data was acquired using an IVIS Spectrum system (Perkin Elmer). The acquired signal was always normalised to radiance (photons/second/centimeter^2^/steradian) and the signal coming from the thoracic area of the animals was quantified using the region of interest (ROI) tool of the IVIS software (Living Image v. 4.5.2) to obtain the total number of photons emitted in that specific area and displayed as total flux (photons/s). Each imaging session was performed using open filter, binning of 8, f-stop of 1-, and 60-seconds exposure time at day 0, and 180 seconds exposure time at days 1, 3 and 7.

#### Statistical analysis

Quantitative data are reported as mean ± standard deviation (SD). N indicates the number of biological replicates, n the number of independent technical replicates. Statistical analyses were performed using GraphPad Prism version 9.2.0 (GraphPad Software, Inc., San Diego, CA, USA). The type of statistical test and the number of replicates included in the analyses are indicated in the figure legends. A p-value < 0.05 was considered statistically significant.

## Results

### Cell culture harmonisation

The first steps to guarantee a reliable head-to-head comparison of the three different MSC sources were directed towards the harmonisation of methodologies across centres. Thus, we defined a common protocol to expand MSCs based on three key parameters: an identical basal medium, namely MEM-α, a batch of FBS and a defined expansion plating density.

Batch-to-batch variability of FBS is a crucial factor in MSC manufacture [21]. We tested three different sera lots on previously isolated BM-MSCs and selected one lot (FBS-A), which promoted growth of MSCs fulfilling the ISCT minimal criteria [22] (**Supplementary Figure 1**).

As plating density can affect proliferation kinetics of MSCs [16, 23], cells from all tissue sources were grown for at least two passages under two seeding densities: 300 and 3,000 cells/cm^2^. At higher seeding density, A- and BM-MSCs had lower cumulative population doublings (CPD), leading to a prolongation of their expansion time (**Supplementary Figure 2a, c, g**). Contrarily, UC-MSCs showed higher CPD when grown at the higher density (**Supplementary Figure 2e**), indicating decreased PDTs (**Supplementary figure 2g**). When assessing cell morphology, UC-MSC lost their spindle-shaped structure when grown at 300 cells/cm^2^ and tended to aggregate and form colonies (**Supplementary Figure 2f**). A similar effect was observed with BM-MSCs, exhibiting a larger and extended cytoplasm (**Supplementary Figure 2d**). The opposite was observed for A-MSCs, which showed a more MSC-like phenotype when grown at 300 cells/cm^2^ (**Supplementary Figure 2b**). Based on these results, BM- and UC-MSCs were expanded at 3,000 cells/cm^2^ while A-MSCs at 300 cells/cm^2^.

### Part A-Biological comparison

In part A of our study, A-, BM- and UC-MSCs, each from three different donors, initiated in one laboratory, were shipped as cryopreserved aliquots to the three sites. Using the harmonised culture protocol (identical FBS lot and culture medium and defined seeding densities), cells were cultured at the three centres for three passages to determine their growth kinetics (**Figure 2a, b**). The results showed that the trends of growth kinetics were consistent across all the sites, despite each type of MSCs being isolated in different laboratories and shipped internationally. BM-MSCs consistently showed the longest PDT in all sites (90.81 ± 10.57 hours - Heidelberg, 66.78 ± 16.32 hours - Galway, 95.72 ± 28.02 hours - Liverpool) as compared to A-MSCs (43.17 ± 3.84 hours, 37.25 ± 1.64 hours, 51.10 ± 1.25 hours in Heidelberg, Galway and Liverpool, respectively) and UC-MSCs (68.07 ± 9.11 hours, 38.06 ± 1.04 hours, 46.06 ± 9.47 hours in Heidelberg, Galway and Liverpool, respectively) (**Figure 2a**). All cells retained their phenotype during culture (**Supplementary Figure 3**). Despite the harmonised culture conditions, some site-to-site variations in PDT were observed (**Figure 2b**), particularly for A- and UC-MSCs where the PDT between sites showed a statistically significant difference. Within all three sites the PDT varied between donors of the same MSC source and between passages of the same donor (**Supplementary Figure 4a-c**). These differences between passages could be observed from the wide distribution of PDTs per donor, as the three data points within a single donor represent PDTs from three consecutive passages. A-MSCs showed the least variation across the different sites and donors. UC-MSCs also showed stable growth throughout the three passages, except in Heidelberg where the difference of PDTs across passages was more prominent than in the other sites. Lastly, BM-MSCs consistently showed high donor-to-donor and passage-to-passage differences in all sites.

**Figure 2.**
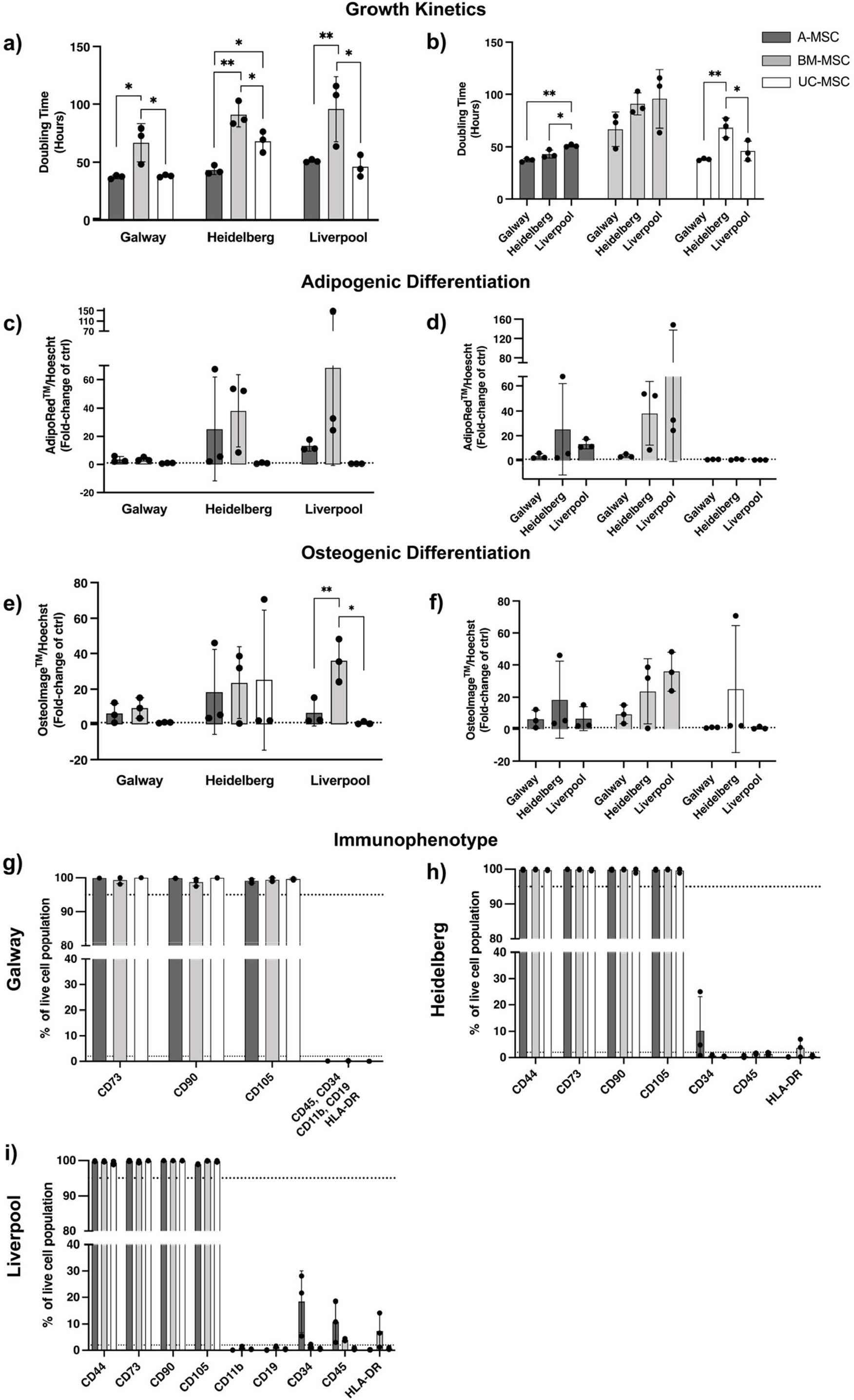
Biological comparison of different tissue sources of MSCs across independent laboratories. (a) In all sites, A- and UC-MSCs showed enhanced growth kinetics when compared to BM-MSCs, with mean doubling times closer to 40 hours for A- and UC-, and 80 to 100 hours for BM-MSCs. (b) Significant differences were observed between sites when comparing the growth rates between sources. (c,d) A- and BM-MSC were able to undergo different levels of adipogenesis and (e,f) osteogenesis while UC-MSCs showed a limited ability to differentiate only into osteocytes (one out of 3 donors at one site). (g-i) Analysis of the immunophenotype by flow cytometry showed adherence to the minimal criteria in all sites, with higher than 95% expression of CD73, CD90 and CD105. Expression of negative markers showed a moderate increase in CD34 (two sites) and CD45 (one site) in A-MSC preparations and a mild increase in HLA-DR in BM-MSC preparations. Data displayed as mean ± SD, N=3. (a) One-Way ANOVA with Tukey’s multiple comparison corrections, * = p < 0.05, ** = p < 0.001, ** = p < 0.001, **** = p < 0.001.

Having established similarities in cell growth, we next assessed the differentiation capacity of the three cell types and between sites (**Figure 2c-f**, data depicted as a fold change of the negative control). Despite the use of harmonised protocols, including commercially available reagents, our results demonstrate high levels of variability, mainly related to inter-lab handling, tissue origin, and donor intrinsic factors. A- and BM-MSCs had a greater propensity to differentiate into adipocytes and osteocytes, despite remarkable differences between sites, while UC-MSCs showed negligible levels of differentiation (**Figure 2c-f**). BM-MSCs displayed the greatest ability to undergo adipogenesis and osteogenesis, but a high degree of variability was observed when comparing inter-lab data and donor-to-donor results (**Supplementary Figure 4d-i**). A-MSC showed similar levels of differentiation in all sites, except one donor showing superior induction abilities in Heidelberg. The wide range of differentiation detected in each site: A- and BM-MSCs possessed considerable higher differentiation abilities for both lineages in Liverpool and Heidelberg. Meanwhile in Galway, differentiation of all MSCs remained relatively modest. Furthermore, greater donor-to-donor variability of MSC differentiation from all tissue sources was more prominent in Liverpool and Heidelberg than in Galway (**Figure 2c-f; Supplementary Figure 4d-i**).

Next, we interrogated the immunophenotype of MSCs using flow cytometry based on the minimal criteria defined previously [22]. Our analysis showed that MSCs from all sources expressed consistently high levels (> 95%) of classical MSC markers CD73, CD90 and CD105 across all sites (**Figure 2g-i**). In Heidelberg and Liverpool, A-MSCs expressed rather high levels of negative surface markers such as CD34 (10.21 ± 12.96% in Heidelberg and 18.40 ± 11.69% in Liverpool) and CD45 (10.81 ± 7.77% in Liverpool). Noticeable levels of HLA-DR were also observed in BM-MSCs in the Heidelberg and Liverpool sites (3.70 ± 3.36% and 7.33± 6.51%, respectively), but not in Galway (**Supplementary figure 4j-l**).

### Part B-Functional *in vitro* comparison

The characterisation of MSCs coming from different sources using the same culture conditions is relatively unexplored and an important step towards defining MSCs in any *in vitro* or *in vivo* comparative study. To investigate whether and to which extent MSCs of different tissue origins differ, we assessed key functional characteristics together with *in vivo* behaviour in part B of this study.

### Angiogenic and endothelial wound healing properties

Support of angiogenesis and endothelial migration is a relevant mechanism of action of MSC-based therapeutics [12]. The angiogenic properties of CM produced by A-, BM-, and UC-MSCs were assessed *in vitro* by testing the ability of their secreted factors to induce endothelial cells to form tubule-like structures when seeded in a Matrigel™ substrate. BM-CM significantly enhanced the generation of a larger and more complex network of tubule-like structures than A- and UC-CM (**Figure 3a**). BM-CM generated tubular networks with significantly more segments (**Figure 3b**), junctions (**Figure 3c**), and closed loops (**Figure 3d**). Evidence of donor-to-donor variability was observed across all cell sources and was statistically significant different for A-MSC – number of junctions – and BM-MSCs – number of junctions and closed loops – (**Supplementary figure 6a-c**). The presence of angiogenic cytokines in MSC-CM was analysed using an antibody array. All sources secreted comparable levels of angiogenic factors; however, differences could be observed in key factors such as VEGF and IGFBP-1 and 2 – higher in BM-CM – or IL-8 and MCP-1 – higher in UC-CM (**Figure 3e**).

**Figure 3.**
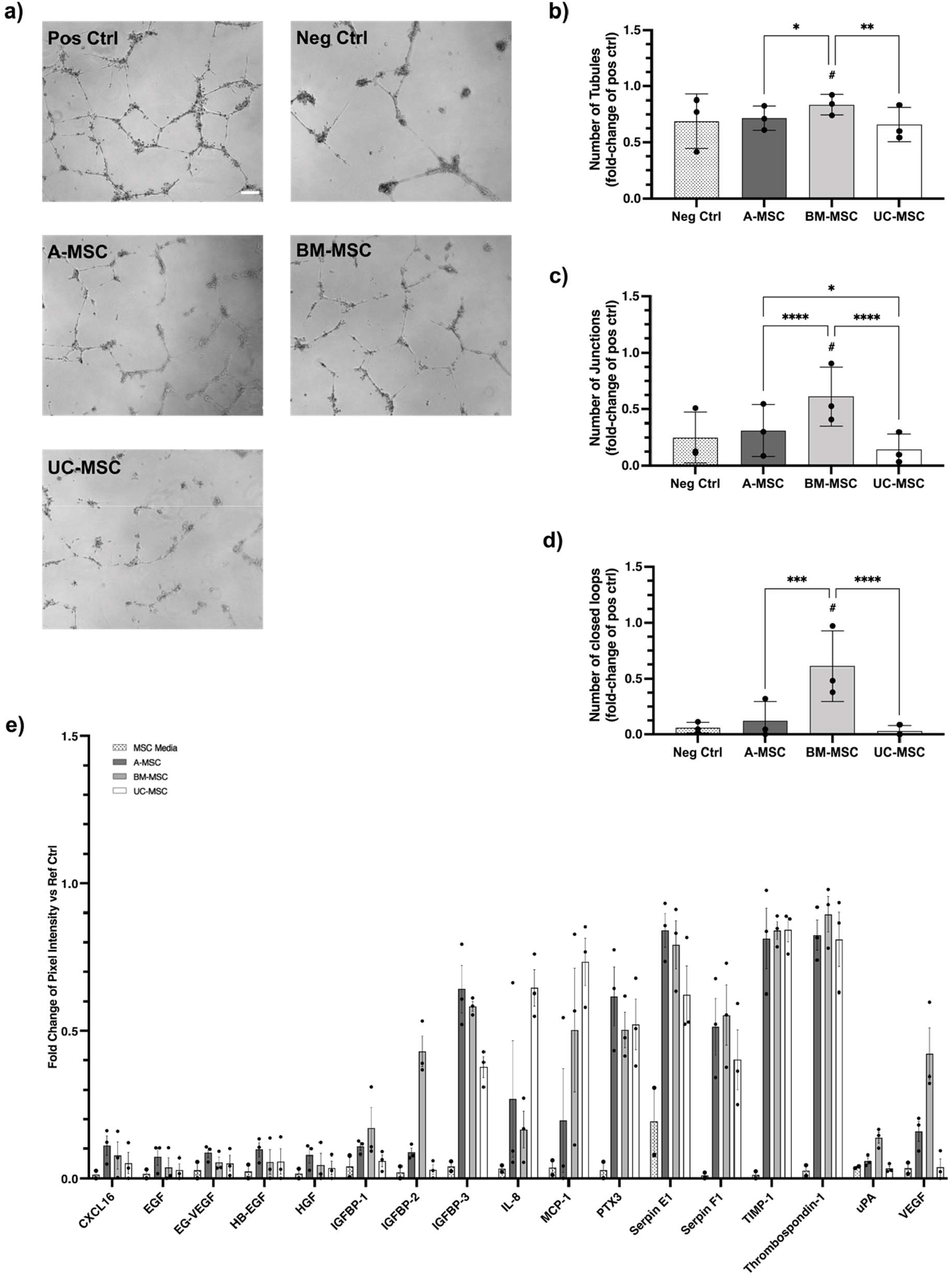
*In vitro* angiogenic properties of MSCs. (a) Representative phase contrast images of tubule-like networks in culture. (b-d) BM-CM generated significantly more tubular-like structures in a more complex and extended mesh (b), represented by a significantly higher number of junctions (c) and closed loops (d) than its counterparts in a model of in vitro tubulogenesis. Data expressed as a fold-change of the positive control. (e) Differential angiogenic proteomic profile for each MSC-CM using an antibody array. Data expressed as a fold change of the reference spots. Data displayed as mean ± SD, N=3, n=3. Two-Way ANOVA with Tukey’s multiple comparison corrections, * = p < 0.05, ** = p < 0.001, ** = p < 0.0001, **** = p < 0.00001. # Significance relative to negative control.

The ability of MSC-CM to induce endothelial cell migration was tested in an *in vitro* wound healing scratch assay. BM-CM resulted in a significant reduction of the scratch gap after 8 and 24 hours (35.03 ± 6.8 % and 58.3 ± 10.36 %, respectively) compared to the negative control (13.73 ± 1.26 % and 3.5 ± 3.3 % at 24 hours; **Figure 4**). The ability of BM-CM to induce endothelial cell migration was significantly superior to A-CM at 24 hours (22.4 ± 2.9 %) and UC-CM at 8 and 24 hours (18.01 ± 1.7 % and 18.1 ± 6.15 respectively) (**Figure 4a**). Limited donor-to-donor variability was observed (**Supplementary figure 6**) although donors with enhanced wound healing properties – such as BM-01 (**Supplementary Figure 6g**) – also exhibited superior abilities to generate tubule-like structures (**Supplementary figure 6a-c**).

**Figure 4.**
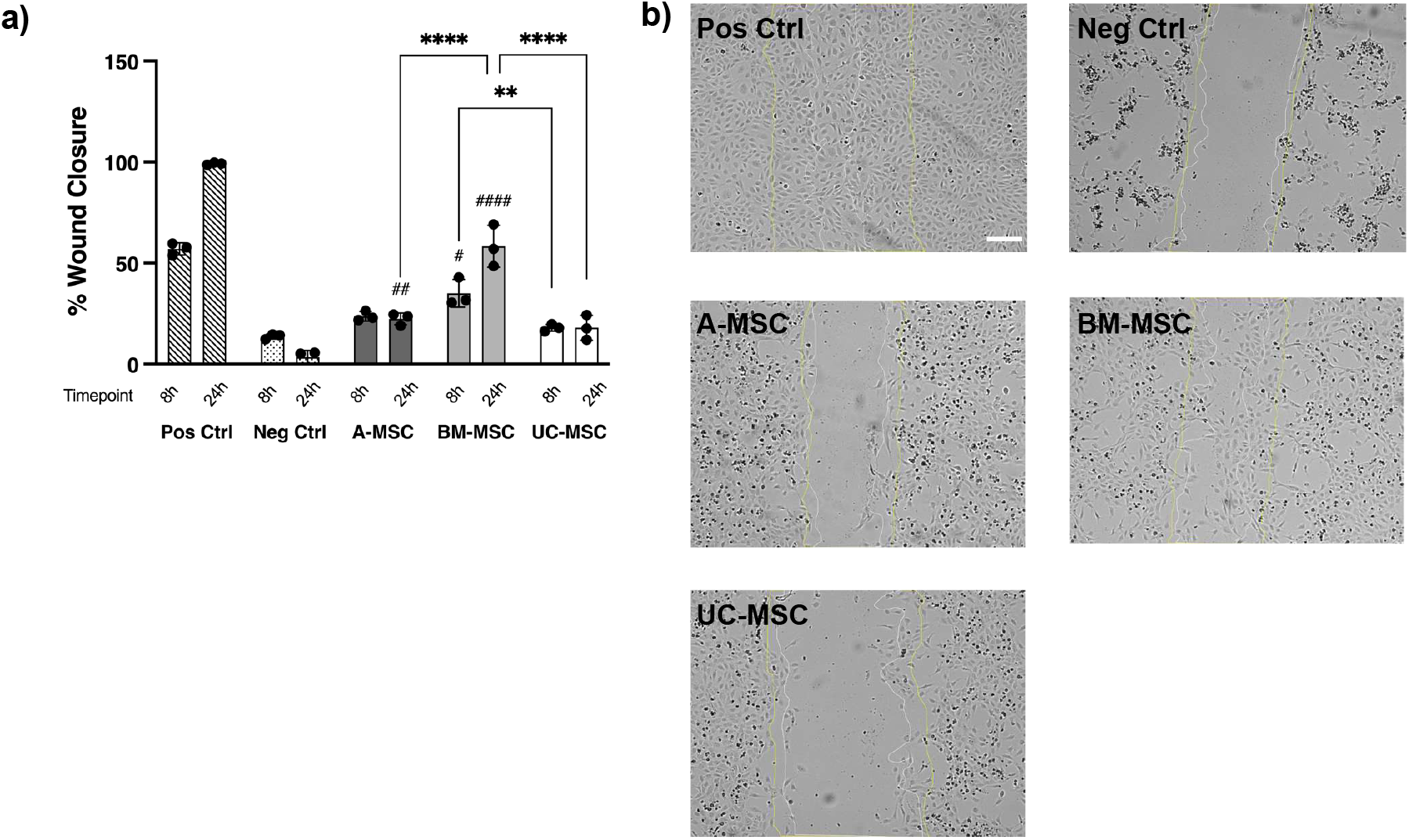
In vitro wound healing properties of MSCs. (a) BM-CM displayed superior ability to induce endothelial cell migration in an in vitro wound healing model at 8 and 24 hours after injury. (b) Representative phase contrast images at time 24 hours after scratch; yellow lines show wound width at time 0 hours and white lines at time 8 hours after scratch. Increased wound gap can be observed at 24h in the negative control due to cell death, when HUVECs are grown with serum-free MEM-α. Data displayed as mean ± SD, N=3, n=3. Two-Way ANOVA with Tukey’s multiple comparison corrections, * = p < 0.05, ** = p < 0.001, ** = p < 0.0001, **** = p < 0.00001. # Significance relative to negative control.

### Immunomodulatory properties

Immunomodulation is a key MSC therapeutic effect [12]. The ability to inhibit PBMC proliferation upon PHA stimulation is often taken as a measure of the immunomodulatory strength [24, 25]. All MSCs were able to suppress PBMC proliferation, as reflected by a decrease in the number of proliferating PBMCs co-cultured with MSCs when compared to those cultured without **(Figure 5a)**. In the presence of A-MSCs PBMC proliferation was significantly reduced (17 ± 0.52 % proliferation relative to positive control), followed by BM-(52 ± 7 %) and UC-MSCs (61 ± 21 %).

**Figure 5.**
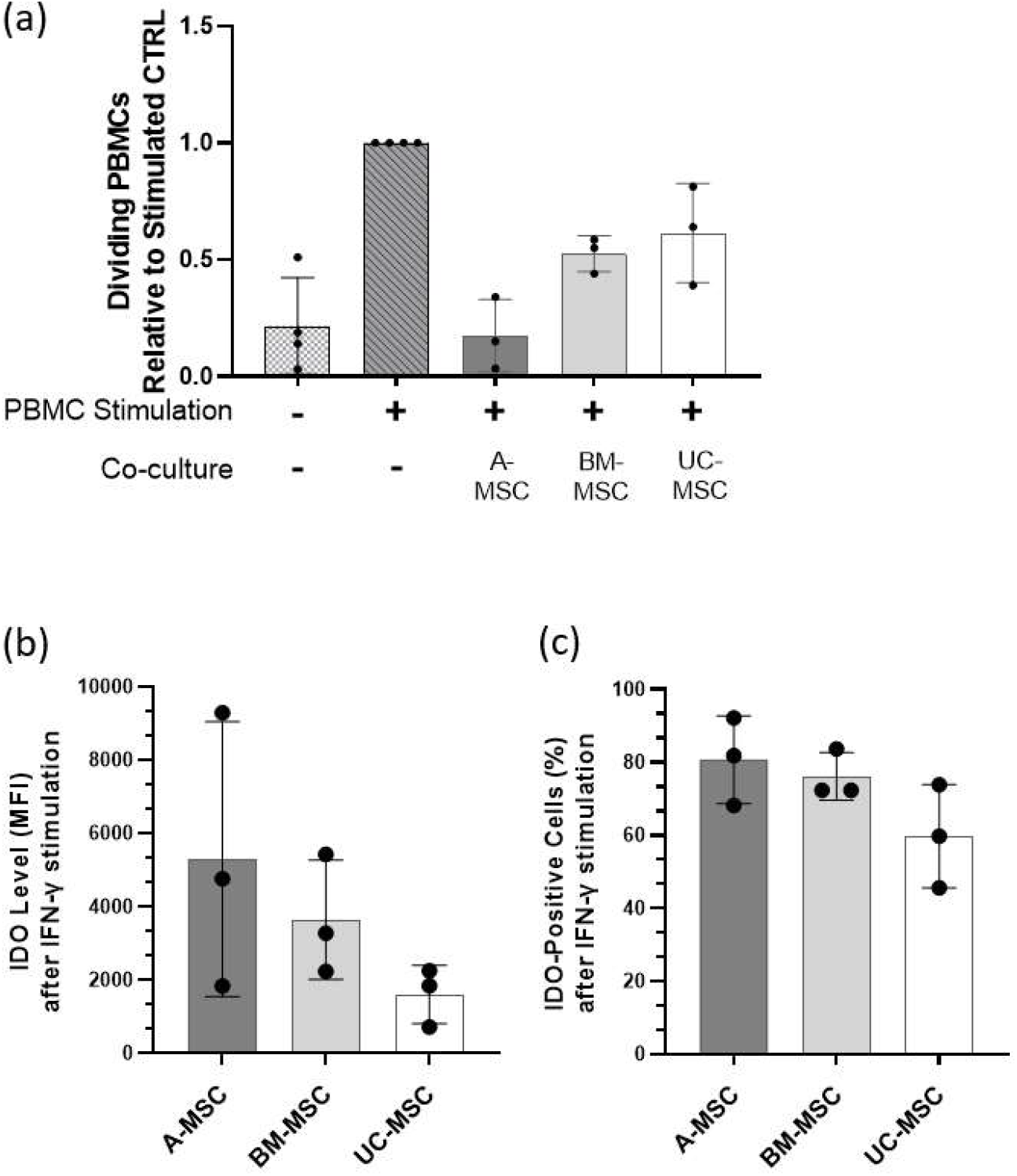
In vitro immunomodulatory capacities of MSCs. (a) PBMC proliferation after five days co-culture with MSCs under PHA stimulation. All values were normalised to PHA-stimulated monoculture PBMCs. (b) Mean fluorescence intensity of intracellular IDO of MSCs after being treated with IFN-γ for 24h. (c) The percentage of cells positive for IDO intracellular staining. Data are displayed as mean ± SD from N=3, n = 3. Two-Way ANOVA with Tukey’s multiple comparison corrections, * = p < 0.05.

The ability of MSCs to inhibit PBMC proliferation was compared to their ability to secrete IDO upon IFN-γ stimulation, since the IDO-kynurenine axis has been shown to be responsible for MSC immunomodulation of T-cells [17]. The level of intracellular IDO, indicated by mean fluorescence intensity (MFI) value, was highest in A-MSCs, followed by BM- and UC-MSCs. High donor-to-donor variability was apparent; highest in A-MSC with values ranging from 5294

± 3752 MFI values (**Figure 5b**). The percentage of cells positive for IDO staining showed the same order, A-MSCs followed by BM- and UC-MSCs; yet here less donor-to-donor variability was observed in all MSC sources (88.77 ± 12.04%, 76.17 ± 6.52% and 59.77 ± 14.15 % for A-, BM-, UC-MSCs, respectively; **Figure 5c)**. Contradicting the notion that IDO levels may correlate with inhibitory strength, donor A-02, the A-MSC donor with the highest ability to suppress PBMC proliferation amongst all A-MSC donors (**Supplementary Figure 7a**), demonstrated the lowest level of intracellular IDO (**Supplementary Figure 7b and c**). In contrast, A-01 with the lowest inhibition of PBMC proliferation amongst A-MSC donors, exhibited the highest level of intracellular IDO.

### *In vivo* biodistribution in healthy mice

We compared the biodistribution of FLuc^+^ MSCs following their IV administration to healthy C57BL/6J albino mice. Regardless of the MSC type, the BLI images reveal that immediately after administration, all signal originating from the injected cells localised to the thoracic region of the body, corresponding to lungs (**Figure 6a**). 24h after infusion the signal was strongly reduced and there was no sign of cell migration from the lungs to any other sites or organs. At this time point, the signal coming from the BM-MSCs seemed weaker than the signal coming from the two other cell types. 3 days after administration a weak signal was detectable from mice that received A- and UC-MSCs, while no signal was detected in most of the mice that received the BM-MSCs. 7 days post administration there was no detectable bioluminescence in any of the animals (**Figure 6a**). These results were confirmed by quantitative analysis of the bioluminescence signal (**Figure 6b and Supplementary Figure 8**). The signal obtained at day 0 was comparable not only between the donors of the same cell type (**Supplementary figure 8**), but also among the different sources of cells (2.8×10^7^ ± 0.99×10^7^ p/s, 4.1×10^7^ ± 0.91×10^7^ p/s and 5.1×10^7^ ± 1.7×10^7^ p/s for A, BM and UC cells respectively; **Figure 6b**). Furthermore, they all showed a similar reduction in the signal from day 0 to day 1 (3.4×10^6^ ± 0.54×10^6^ p/s, 0.83×10^6^ ± 0.9×10^6^ p/s and 3.6×10^6^ ± 2.5×10^6^ p/s for A, BM and UC cells respectively) and to day 3 (8.3×10^5^ ± 2.0×10^5^ p/s, 1.6×10^5^ ± 076×10^5^ p/s and 2.9×10^5^ ± 1.1×10^5^ p/s). By day 7 the detected signal (1.04×10^5^ ± 0.11×10^5^ p/s, 1.07×10^5^ ± 0.26×10^5^ p/s and 0.97×10^5^ ± 0.09×10^5^ p/s respectively) was no different from the naïve animals (1.1×10^5^ ± 0.07×10^5^ p/s) that did not receive any cells or substrate. The analysis of relative bioluminescence intensity normalised to signal at day 0 revealed that in the first 24 hours the signal dropped significantly to 12.9 ± 3.4% for the A-MSCs, to 2.5 ± 3.1% for the BM cells and to 6.3 ± 3.6% for the UC cells (**Figure 6c**).By day 3, only 3.47 ± 1.7%, 0.44 ± 0.31% and 0.58 ± 0.05% of the original signal was detectable for A-, BM- and UC-MSCs, respectively.

**Figure 6.**
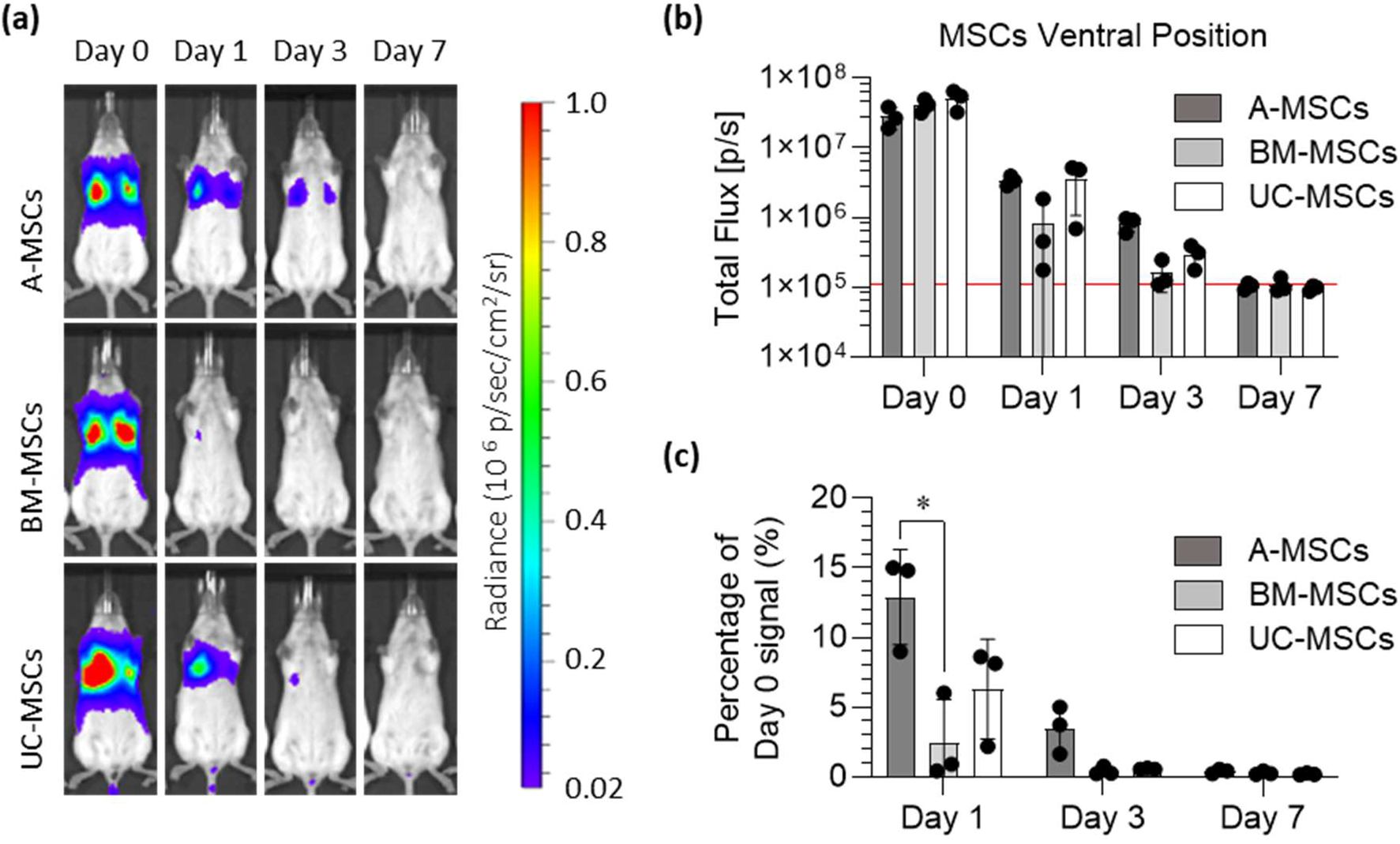
All MSCs were entrapped in the lungs and were short-lived following IV administration. (a) Representative bioluminescence images of mice administered with FLuc expressing A-, BM- and UC-MSCs on the day of administration of the cells (day 0), and after 1, 3 and 7 days (radiance scale from 0.2×10^5^ to 1×10^6^ p/s/cm^2^/sr). (b) Light output (flux) as a function of time (days) from the three different types of MSC. (c) Signal at day 1, day 3, and day 7 normalised to day 0 signal. Data in charts are displayed as mean ± SD from three donors for each type of MSC (4 animals used per donor). Two-Way ANOVA with Tukey’s multiple comparison corrections, * = p < 0.05.

## Discussion

Within this study, we first aimed to assess the impact of different decentralised production sites on MSC characteristics and second to understand differences in tissue source specific properties.

Contrary to Stroncek *et al*., who shipped the same tissue starting material to the different manufacturing sites [26], we mimicked the situation of one initial manufacturing centre and different decentralised cell production facilities that expand MSCs using harmonised protocols and quality control the final MSC product. We pre-defined harmonised conditions by culturing all three MSC types in the same MEM-α supplemented with the same lot of FBS. Finally, to properly compare the different sources, a seeding density optimal for the expansion of each cell type was identified and adopted across centres.

In part A, our study shows for the first time that the protocol harmonisation reduces to some extent site-to-site variation whilst the tissue and donor-specific differences remain apparent. BM-MSCs exhibited the longest doubling time as well as the highest inter-donor variability, whereas A-MSCs consistently showed the least donor-to-donor variation regardless of where they were cultured. Site-to-site variation can in part be attributed to the differing shipment duration on dry ice, which interrupted the cold chain. The manual handling of cell counting and assessment of confluence for harvest also contributed to the site-to-site variations. Given that MSCs show contact-dependent growth inhibition [23], slight differences in the confluence may affect the calculation of growth kinetics. More objectified, operator-independent, assessment of confluence and cell counting is expected to significantly improve comparability.

The analysis of adipogenic and osteogenic potential confirmed the known inter-donor variability that was consistent in all sites. Despite the use of harmonised differentiation protocols and kits, quantitative results varied largely, demonstrating the large influence exerted by the operator. UC cells displayed no adipogenic or osteogenic potential in any of the centres. Reduced or entire lack of adipogenic differentiation potential has been repeatedly reported for perinatal MSCs [27, 28]. Yet, the entire lack of *in vitro* osteogenic differentiation in UC-MSCs (with one most probably artefactual outlier in one site) was rather unexpected. It may reflect differing requirements of UC-MSCs for osteoinduction [29]. However, it is not clear whether the *in vitro* differentiation potential is a meaningful selection criterion when defining the best source of MSC for the intended therapeutic application [4, 10]. We suggest that if differentiation potential is taken as critical attribute, it should be assessed qualitatively, or if quantitatively, as a batch comparison within one centre.

Expression of surface markers (including CD73, CD90 and CD105) and lack of hematopoietic markers (including CD11b, CD19, CD34 and CD45) and major histocompatibility complex (MHC) class II (HLA-DR) are widely accepted criteria to assess the identity and purity of MSCs [22]. Whilst in the three centres MSCs from all donors showed a positivity of at least 98% for all the positive markers, some variability was observed for the negative ones. In particular, A-MSCs showed increased expression (> 2%) of CD34 (Heidelberg and Liverpool) and CD45 (Liverpool). This is not unexpected as previous studies have reported CD34 positivity of A-MSCs, at least early in culture [30-32]. Similar early expression of CD45 disappearing after prolonged culture was also observed in BM-MSCs [33]. Moreover, 2 of the 3 BM-MSCs showed a small variability in the positivity to HLA-DR in two centres (Heidelberg and Liverpool). Similar findings have been previously reported by Grau-Vorster *et al*. who revealed variability in BM-MSC preparations for clinical applications, concluding that the absence or presence of HLA-DR does not have an impact on the overall properties of the cells [34]. Of note, CD34 and HLA-DR positivity observed in the two separate sites in the same donors, strongly suggests donor-related variability as the main cause.

It is noteworthy that in this study, the cells were isolated in one specific centre, cryopreserved, and then shipped in dry ice before being expanded and compared in each site in parallel. Cryopreservation not only affects the proliferation of the cells [35], but also impacts the differentiation potential [36] and the immunosuppressive properties [37]. However, it has been described to be a transient effect due to the heat-shock stress induced by the thawing process, with functionality being restored after a certain culture period [38]. In this study, the effect of international shipping has not been evaluated in detail. Our data and that of Stroncek, however, clearly suggest that before such a study, cultivation and quality control protocols require not only harmonisation but rather standardisation to minimise site-specific influences as much as possible.

To determine whether the heterogeneity of MSCs from different origins is also reflected in their potential therapeutic abilities, part B of our study provided a comparison of the tissue sources on top of basic cell characteristic assessments. This comparison was performed each in a single expert centre. First, we assessed the angiogenic profile of CM obtained from A-, BM- and UC-MSCs. In our hands, CM from BM-MSCs showed superior abilities to form tubule-like structures and induce endothelial cell migration *in vitro*. The overall presence and concentration of angiogenic factors within the CM was found to be superior in BM preparations with increased relative levels of tubulogenesis-driving factors such as VEGF [39, 40]. Although our results align with previous studies showing superior proangiogenic abilities [41] and higher secretion of VEGF in BM-MSC cultures [42], others have conversely reported higher tube formation and angiogenic bioactivity in the secretome of A-MSCs [43, 44]. Most likely, technical discrepancies along with donor-to-donor variability are playing key roles. For instance, dose-dependent levels of VEGF from BM-MSC secretomes have been correlated with angiogenic activity and proposed as a surrogate potency assay for clinical preparations [45]. Donor variability is a well-known phenomenon we have also observed within our sample preparations, emphasising the need to dissect donor characteristics and variability in autologous and allogeneic settings to achieve favourable clinical outcomes [46].

Second, we investigated whether the source of MSCs might influence their immunomodulatory capacity to suppress PBMC proliferation. We also measured IDO production after IFN-γ stimulation as IDO has been implicated as the key factor responsible for inhibition of PBMC proliferation by catabolism of tryptophan to kynurenine [17, 47]. A-MSCs, the tissue source of MSCs with the highest ability to inhibit PBMC proliferation, exhibit the highest level of intracellular IDO upon IFN-γ stimulation, followed by BM and UC-MSCs. Our data however question a correlation between IDO levels and proliferation inhibitory strength, given that the donor which showed the highest inhibition exhibited the least intracellular IDO and vice versa. Although we previously showed that MSC-expressed IDO is key to inhibit PHA-driven T cell proliferation [17], this is most likely not the only factor involved, especially when considering the much more complex situation in vivo. A study by Chinnadurai *et al*. elegantly showed that MSCs can inhibit PBMC proliferation through PD1/PD-L1 [48].

Our data demonstrate that the different MSC types have individual properties, which may have benefits in specific therapeutic settings. A-MSC show enhanced immunoregulatory abilities, BM-MSC superior angiogenic and wound healing properties while UC-MSC appears to be the least potent of all three sources. In this sense, whether the assays proposed are able to capture all the properties and attributes from each tissue source needs further validation in specific *in vitro* and *in vivo* injury models to confirm their ability to predict therapeutic potency. A more detailed and complex picture of their secretome, including the shedding of extracellular vesicles [49] and microRNAs [50], the mitochondrial and metabolic properties [51], together with other aspects of their immunomodulatory properties not addressed in this study, might highlight further attributes aligned with desirable clinical outcomes.

Third, an important aspect of this study was to investigate and compare the fate of different MSCs *in vivo* after being cultured using the same manufacturing procedures. Intravenous administration of MSCs is the most common delivery route used in clinical trials [52]. However, it is well known that MSCs get entrapped in the lung, the so called pulmonary first pass effect [53-55]. Besides posing a risk for embolisation, pulmonary trap reduces the number of cells that could eventually home and engraft to the injured tissue [56]. Here, BLI performed immediately after the IV administration of different MSCs in healthy mice confirmed their entrapment in the lungs, irrespective of their tissue of origin. Additionally, none of the cells escaped the lungs, neither on the day of administration nor in any of the following days. In fact, a major drop in the bioluminescence signal coming from the lungs was observed in the first 24h post injection. Despite signal from A-MSCs being still noticeable 3 days post administration, no signal from any of the MSCs was detected 7 days after injection. This result is consistent with various reports [54, 55], and confirms that this effect is not influenced by MSC origin. When cell therapies are considered, the fact that most of MSCs die in the first 24 hours is not necessarily a bad result. It has been proposed that the apoptosis of IV administered MSCs in the lungs and the subsequent phagocytosis of the cell debris by local macrophages is a mechanism of MSC-mediated immunomodulation [55, 57-59].

In summary, we have:

## Conclusions

Lack of standard culture protocols is a major limitation that hinders comparison of the clinical benefits of MSCs, especially when they are from different sources, and produced in different centres. Here we established, for the first time, harmonised tissue culture conditions for expansion of A-, BM- and UC-MSCs among three independent centres across Europe to investigate the reproducibility of these procedures and its impact on their biological characteristics and functionality both *in vitro* and *in vivo*. We show that harmonised protocols improve reproducibility across different centres emphasising the need for worldwide standards to manufacture MSCs for clinical use. Further, tissue-specific differences in cell characteristics suggest a need for selecting the optimal cell type for the intended clinical indication based on source availability and functional characteristics. These results show the heterogeneous behaviour and therapeutic properties of MSCs as a reflection of tissue-origin properties while providing evidence that the use of harmonised culture procedures can reduce but not eliminate inter-lab and operator differences.

## Author Contributions

All authors have made substantial contributions to all of the following: (1) the conception and design of the study, or acquisition of data, or analysis and interpretation of data, (2) drafting the article or revising it critically for important intellectual content, (3) final approval of the submitted version.

## Acknowledgments

The team in Galway would like to acknowledge technical and consultative support for flow cytometry experiments provided by Dr. Shirley Hanley of the University of Galway Flow Cytometry Core Facility, which is supported by funds from University of Galway, Science Foundation Ireland, the Irish Government’s Programme for Research in Third Level Institutions, Cycle 5 and the European Regional Development Fund.

The team in Mannheim acknowledges the excellent support of Stefanie Uhlig, FlowCore Mannheim and Institute of Transfusion Medicine and Immunology, and Corinna Thielemann for excellent technical support.

The team in Liverpool acknowledges the support of the Flow Cytometry Facility.

## Abbreviations

A: Adipose
BLI: Bioluminescence Imaging
BM: Bone Marrow
CM: conditioned Media
CPD: Cumulative Population Doublings
DEAE-dextran: Diethylaminoethyl-Dextran
DMSO: Dimethyl Sulfoxide
DPBS: Dulbecco’s phosphate-buffered saline
EGM: Endothelial Growth Medium
EU: European Union
FACSs: Fluorescence Activated Cell Sorting
FBS: Foetal Bovine Serum
HEK: Human Embryonic Kidney cells
HLA-DR: Human Leukocyte Antigen – DR Isotype
HUVEC: Human Umbilical Cord Endothelial Vein Cells
IDO: Indoleamine 2,3-dioxagenase
IFN-: γ Interferon gamma
IGFBP: Insulin-like Growth Factor Binding Protein
IL: Interleukin
ISCT: International Society for Cell and Gene Therapy
ITN: Innovative Training Network
IV: Intravenously
LV: Lentiviral Vector
MCP-1: Monocyte Chemoattractant Protein 1
MEM-: α Minimum Essential Medium Alpha
MFI: Mean Fluorescence Intensity
MHC: Major Histocompatibility Complex
MOI: Multiplicity of Infection
MSC: Mesenchymal Stromal Cell
PBMC: Peripheral Blood Mononuclear Cells
PD: Population Doublings
PDT: Population Doubling Time
PHA: Phytohemagglutinin-L
ROI: Region of Interest
RPMI: Roswell Park Memorial Institute Medium
SC: Subcutaneous
SD: Standard Deviation
UC: Umbilical Cord
VEGF: Vascular Endothelial Growth Factor

## Supplemental Information

### Materials and Methods

#### 1. Serum Screen

Three batches of FBS from different suppliers were tested to source a serum that supported the growth of MSCs from each tissue source in adequate amounts to service the complete project. Three already isolated BM-MSC donors were used, and we measured their proliferation rates, immunophenotype, and trilineage differentiation potential. Proliferation and immunophenotype were performed as described previously in the Materials and Methods section.

For adipogenic differentiation, confluent cultures were treated with adipogenic induction media, which consisted of high glucose Dulbecco’s modified eagle medium (HG-DMEM, SigmaAldrich, D5796) supplemented with 10% of each FBS respectively and 1% penicillin-streptomycin (PS, Gibco, 15140-122), 1 µM dexamethasone (Merck, D4902), 10 µg/mL insulin (Sigma, 11376497001), 200 µM indomethacin (Merck, I7378) and 500 µM 3-Isobutyl-1-Methyl-Xanthine (MIX, Merck, I7018)) during 3 days and subsequently with adipogenic maintenance media (HG-DMEM, 10% of each FBS, and 1% PS) for 1 day in three repeating cycles. At the completion of the last cycle, cells were incubated in maintenance media for 7 days. Control cells were maintained in regular culture media. Detection of intracellular lipid accumulation was achieved by staining the cultures with 3% Oil Red O (Sigma, O0625) in de-ionised water (6:4) after fixation with 10% neutral buffered formalin (Sigma-Aldrich, HT501128). Harris Modified Haematoxylin (Sigma-Aldrich, HHS-16) was used to counterstain before brightfield imaging at 4X on an Olympus BX43 microscope fitted with an HD Chrome camera (1/.8”) and a 0.5x C-mount adapter. For quantitative analysis, 99% isopropanol (Sigma-Aldrich, I9516) was used to extract the Oil Red O-stained lipids that were then quantified in a multimode plate reader via absorbance at 490 nm (Victor X3, Perkin Elmer).

For chondrogenic differentiation, harvested BM-MSCs were transferred to screw capped microcentrifuge tubes at a concentration of 2 × 10^5^ cells and centrifuged at 100 g for 5 minutes in a swing out rotor to generate cell pellets. Negative differentiated pellets were cultured with incomplete chondrogenic media (ICM: HG-DMEM supplemented with 100 nM dexamethasone, 50 µg/mL ascorbic acid 2-phosphate (Sigma, A8960) 40 µg/mL L-proline (Sigma, P0380), 1X ITS+ media supplement (insulin, transferrin, selenous acid, linoleic acid, bovine serum albumin), 1 mM sodium pyruvate,1% PS) while positive differentiated pellets were cultured with complete chondrogenic media (CCM: ICM supplemented with 10 ng/ml transforming growth factor β3 (TGF-β3, R&D Systems, UK). Media changes were performed every other day for 21 days. The level of chondrogenesis was assessed by measuring the sulphated glycosaminoglycans (s-GAG) present in each pellet using the DMMB assay and normalising between cultures by DNA content measured using PicoGreen™ Quant-iT Kit (ThermoFisher Scientific, P11496) as per the manufacturer’s instructions.

For osteogenic differentiation, 80% confluent cultures were treated with osteogenic induction media, which consisted oflow glucose Dulbecco’s modified eagle medium (LG-DMEM, SigmaAldrich, D6046) supplemented with 10% of each FBS, 1% PS, 100 nM dexamethasone, 100 µM ascorbic acid 2-phosphate (Sigma, A8960), and 10 mM ß-glycerophosphate (Sigma, G9422). Control cells were maintained in regular culture media. Media changes were performed twice per week for 17 days. At the end of this period, cultures were washed with PBS and treated with 0.5 M HCl (Sigma, 1090581000) to collect the cell layer. The cell suspension was then incubated overnight at 4 °C under agitation, centrifuged to discard cell debris and the calcium present in the supernatant quantified using the Stanbio Calcium CPC liquicolour kit (Stanbio via ThermoFisher, 0150250) as per the manufacturer’s instructions. Representative brightfield images of the cultures were taken after fixation in 95% ice cold methanol and staining for calcium deposits with 2% Alizarin Red S (Merck, A5533). Images were taken as described for adipogenic cultures.

## Supplementary Tables

**Supplementary Table 1.**
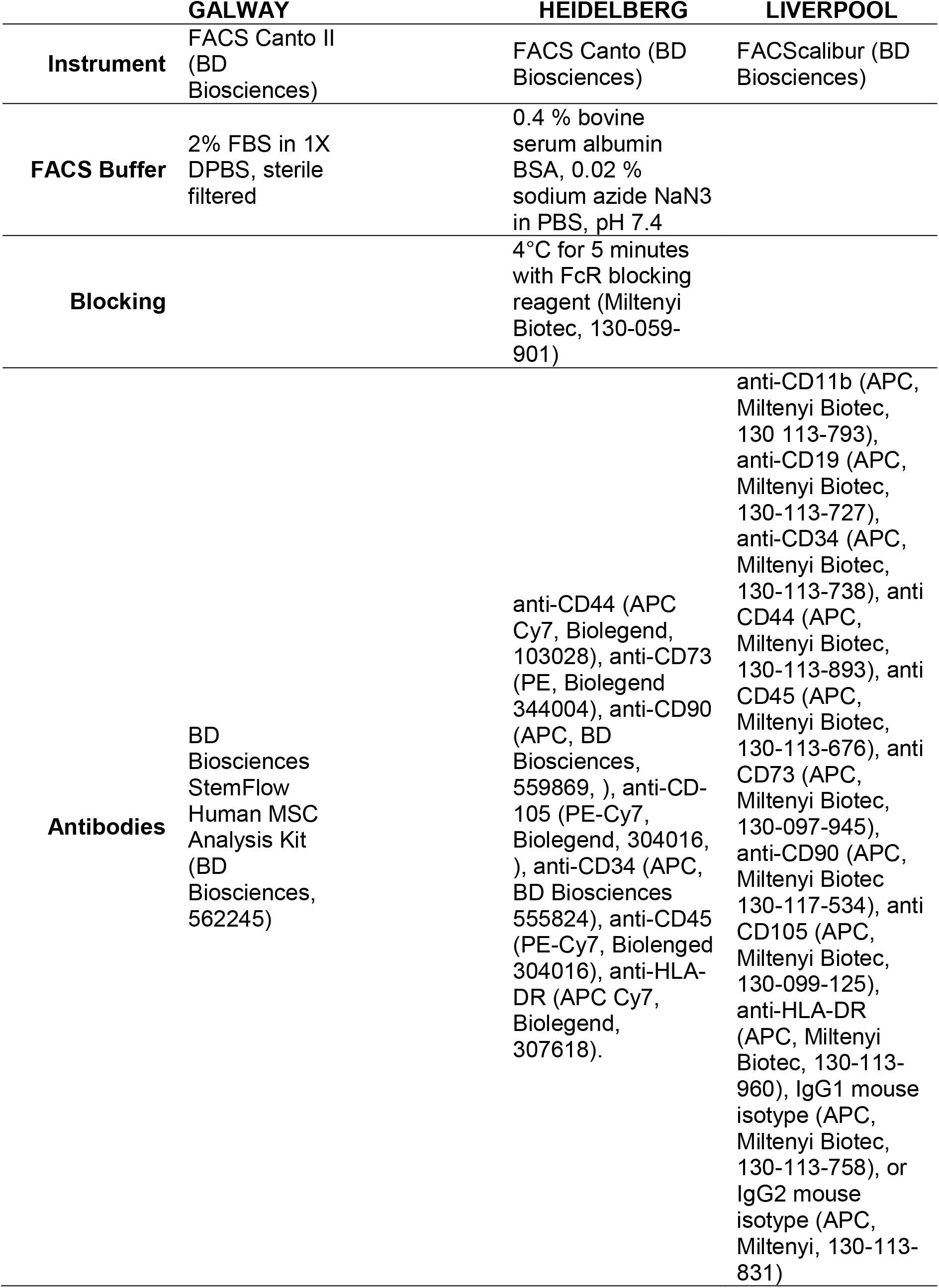
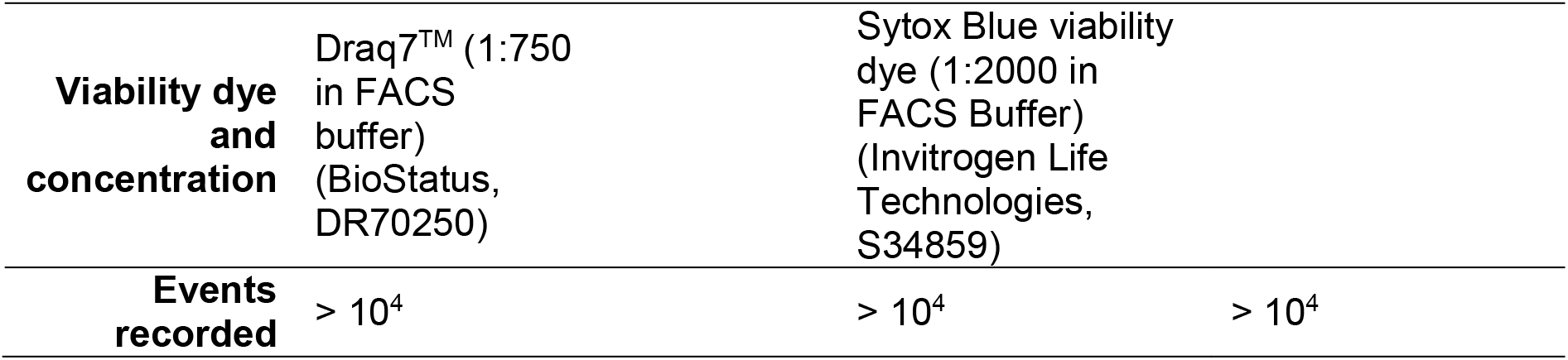
Detailed methodology used in each centre to characterise the immunophenotype of MSCs

|  | **GALWAY** | **HEIDELBERG** | **LIVERPOOL** |
| --- | --- | --- | --- |
| **Instrument** | FACS Canto II<br>(BD Biosciences) | FACS Canto (BD Biosciences) | FACScalibur (BD Biosciences) |
| **FACS Buffer** | 2% FBS in 1X DPBS, sterile filtered | 0.4 % bovine serum albumin BSA, 0.02 % sodium azide NaN<sub>3</sub> in PBS, pH 7.4 |  |
| **Blocking** |  | 4°C for 5 minutes with FcR blocking reagent (Miltenyi Biotec, 130-059-901) |  |
| **Antibodies** | BD Biosciences StemFlow Human MSC Analysis Kit (BD Biosciences, 562245) | anti-CD44 (APC Cy7, Biolegend, 103028), anti-CD73 (PE, Biolegend 344004), anti-CD90 (APC, BD Biosciences, 559869, ), anti-CD-105 (PE-Cy7, Biolegend, 304016, ), anti-CD34 (APC, BD Biosciences 555824), anti-CD45 (PE-Cy7, Biolenged 304016), anti-HLA-DR (APC Cy7, Biolegend, 307618). | anti-CD11b (APC, Miltenyi Biotec, 130-113-793), anti-CD19 (APC, Miltenyi Biotec, 130-113-727), anti-CD34 (APC, Miltenyi Biotec, 130-113-738), anti CD44 (APC, Miltenyi Biotec, 130-113-893), anti CD45 (APC, Miltenyi Biotec, 130-113-676), anti CD73 (APC, Miltenyi Biotec, 130-097-945), anti-CD90 (APC, Miltenyi Biotec 130-117-534), anti CD105 (APC, Miltenyi Biotec, 130-099-125), anti-HLA-DR (APC, Miltenyi Biotec, 130-113-960), IgG1 mouse isotype (APC, Miltenyi Biotec, 130-113-758), or IgG2 mouse isotype (APC, Miltenyi, 130-113-831) |
| **Viability dye and concentration** | Draq7™ (1:750 in FACS buffer) (BioStatus, DR70250) | Sytox Blue viability dye (1:2000 in FACS Buffer) (Invitrogen Life Technologies, S34859) |  |
| **Events recorded** | > 10<sup>4</sup> | > 10<sup>4</sup> | > 10<sup>4</sup> |

## Supplementary Figures

**Supplementary Figure 1:**
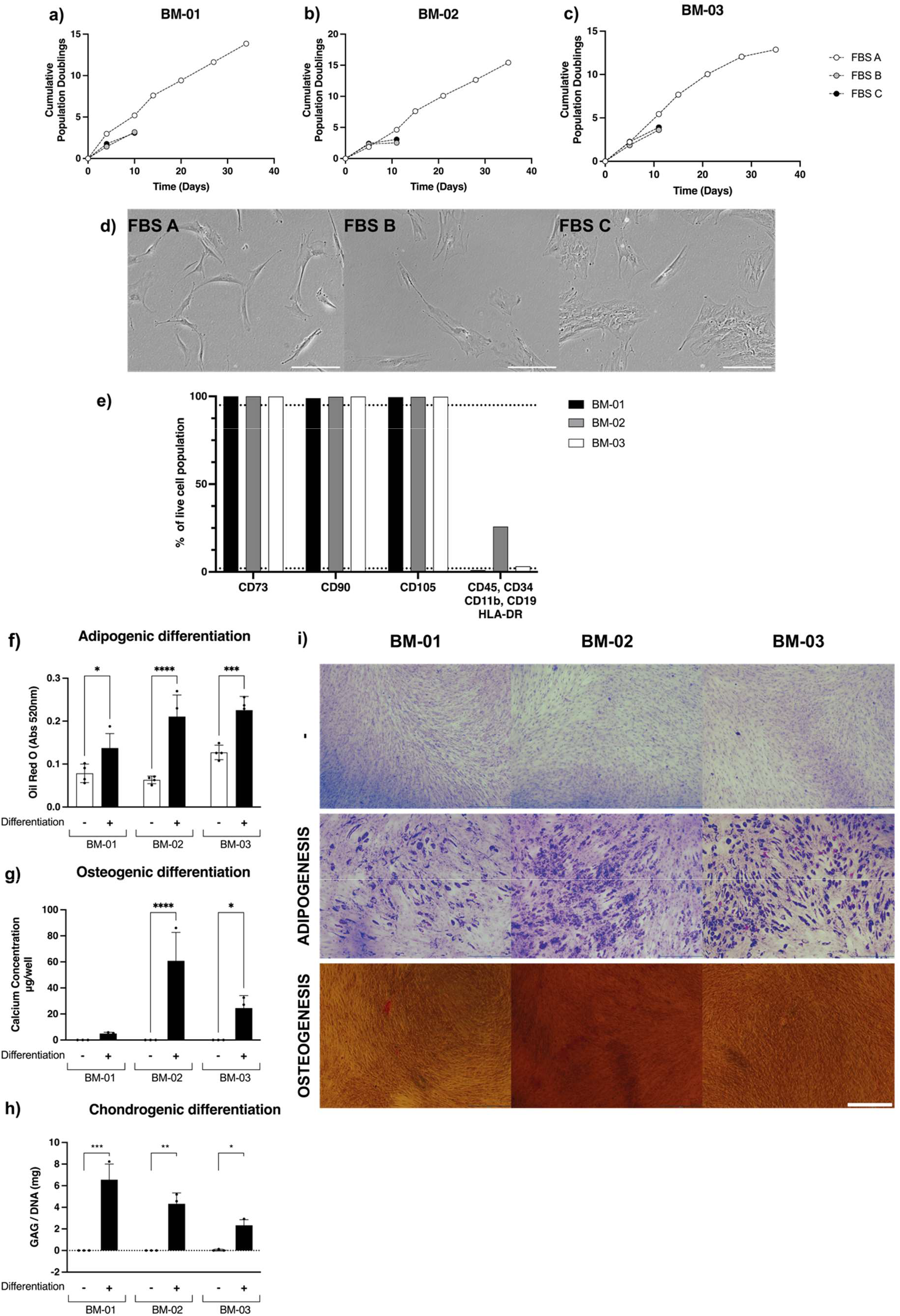
Cell Culture Harmonisation: Serum Screen. (a-c) Population doubling times and (d) phase contrast images of three BM-MSCs donors showed that exclusively FBS A supported cell growth and fibroblast-like morphology. Therefore, further experiments were carried out using serum A. (e) Flow cytometry confirmed the expression of positive surface antigens (CD90, CD73, CD105) and lack of negative markers (CD45, CD34, CD11b, CD19, HLA-DR) in two out of three populations grown with FBS A. (f-i) BM-MSC cultures were induced to differentiate into adipocytes (+) while undifferentiated cultures served as control (-). Images of Oil Red O are shown in panel (i) and quantification of Oil Red O stain retention in panel (f). Both show an increase in lipid content in the majority of adipogenic differentiated cultures. (i) Osteogenic differentiated cultures showed presence of calcium in the extracellular matrix with Alizarin Red staining. (g) Quantification of extracted calcium from osteogenically differentiated BM-MSC showed more than 1 µg of calcium per well in all differentiated cultures. (h) Quantification of sulphated glycosaminoglycans (s-GAG) showed significantly increased levels in differentiated cultures (+), confirming their mesodermal differentiation abilities. Data displayed as mean ± SD, N=3. Two-Way ANOVA with Bonferroni’s multiple comparison corrections, * = p < 0.05, ** = p < 0.001, ** = p < 0.0001, **** = p < 0.00001. Pictures taken at 40X; scale bar 500 µm.

**Supplementary Figure 2.**
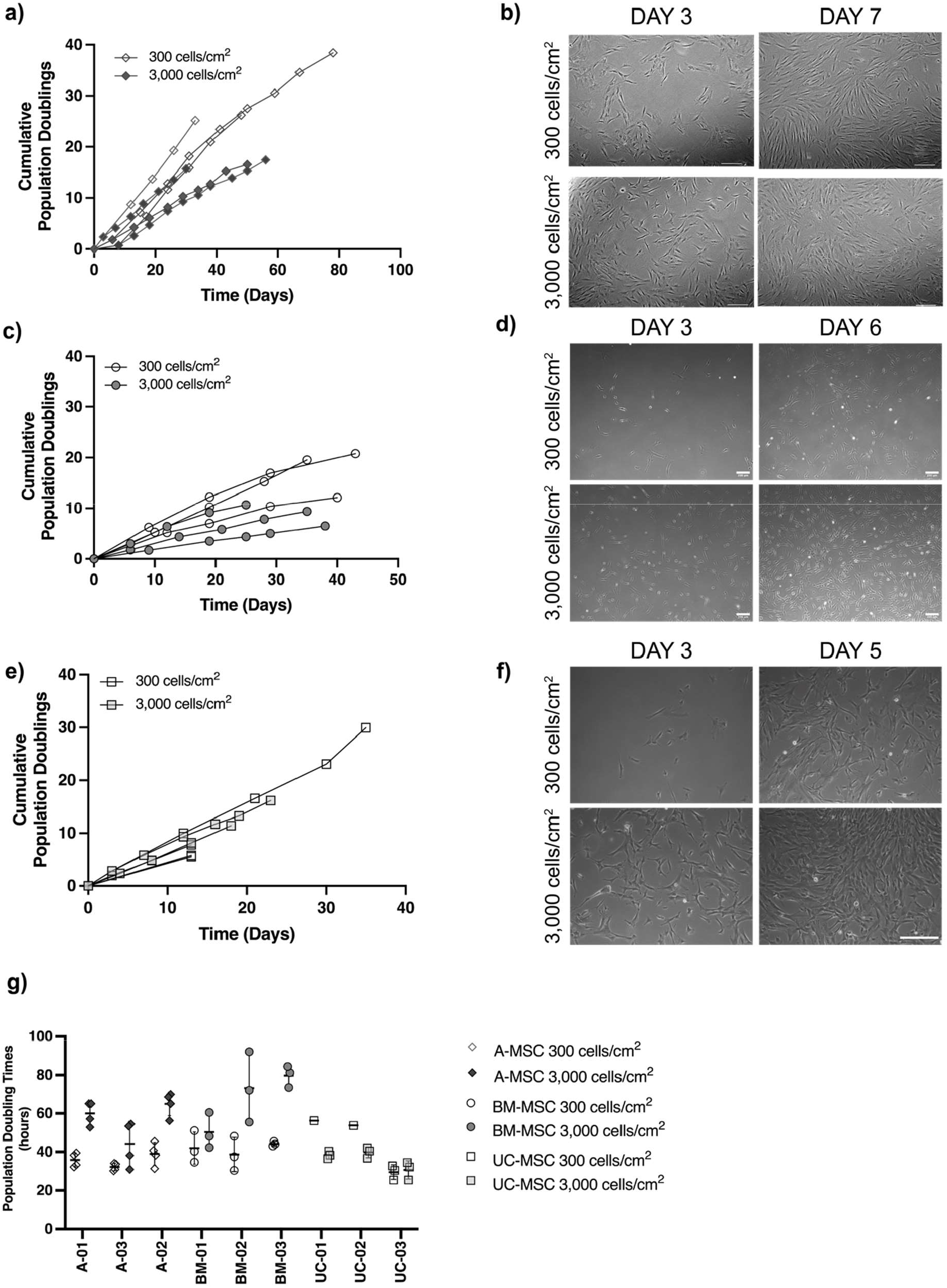
Cell Culture Harmonisation: Seeding Density. Comparison between seeding density confirmed differences in cell source. Cumulative population doublings were calculated by culturing MSCs at 300 (empty symbols) and 3,000 cells/cm^2^ (filled symbols) in all three different sites. (a) A-MSC and (c) BM-MSC showed a rapid increase in cumulative doublings when seeded at a lower density versus at high density after the same period in culture. (e) UC-MSC conversely had increased cumulative doublings when seeded at higher density. (g) When comparing population doubling times, A-MSC and BM-MSC had prolonged kinetics when grown at 3,000 cells/cm^2^ whereas UC-MSC divided faster at 3,000 cells/cm^2^. (b,d,f) Representative phase contrast images of MSCs. Data displayed as mean ± SD, N=3. Pictures taken at 40X.

**Supplementary Figure 3.**
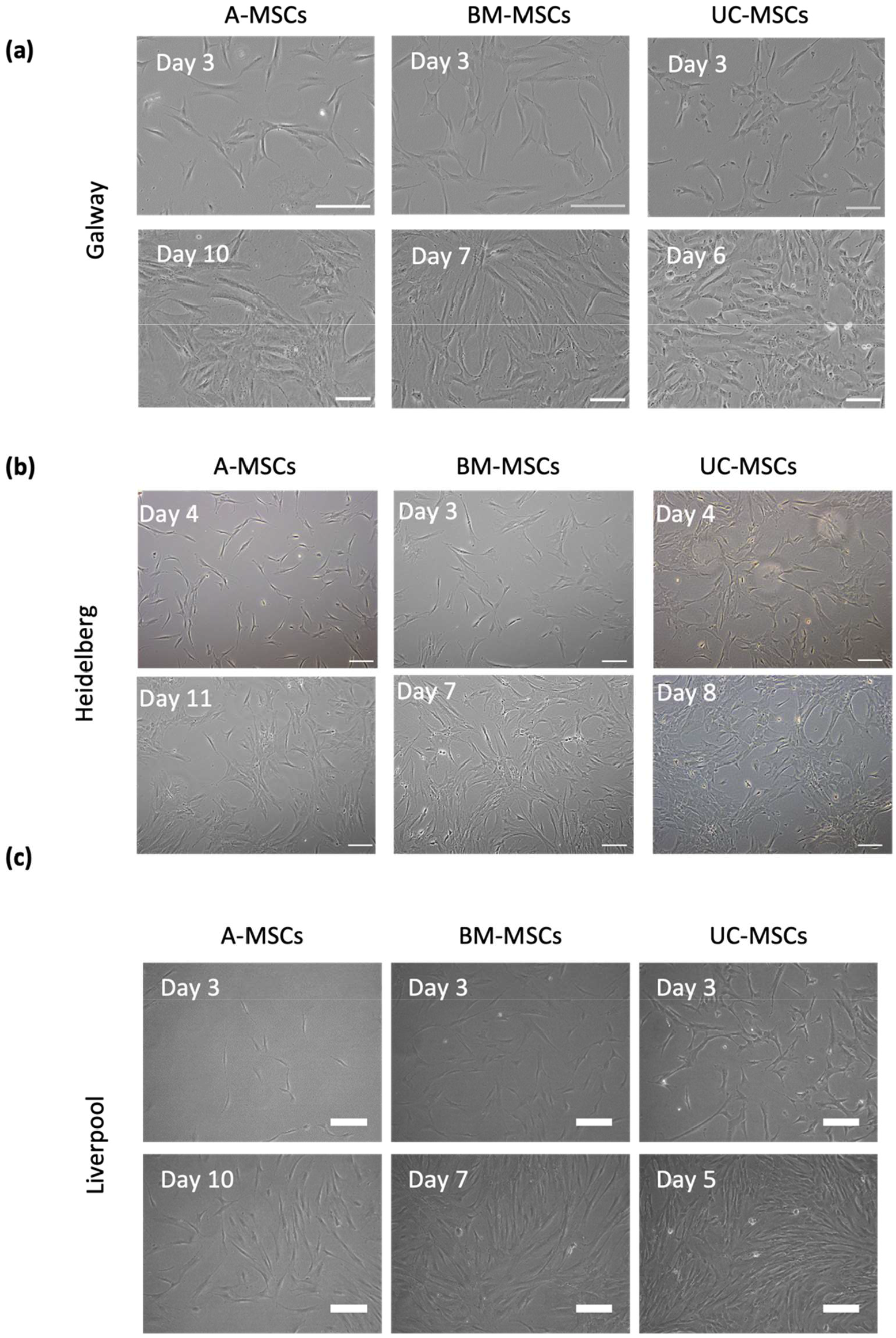
Representative phase contrast images of MSCs in all sites at early (3-4 days) and late (5-10 days) stages of culture. Pictures taken at 100X; scale bar 200 µm.

**Supplementary Figure 4.**
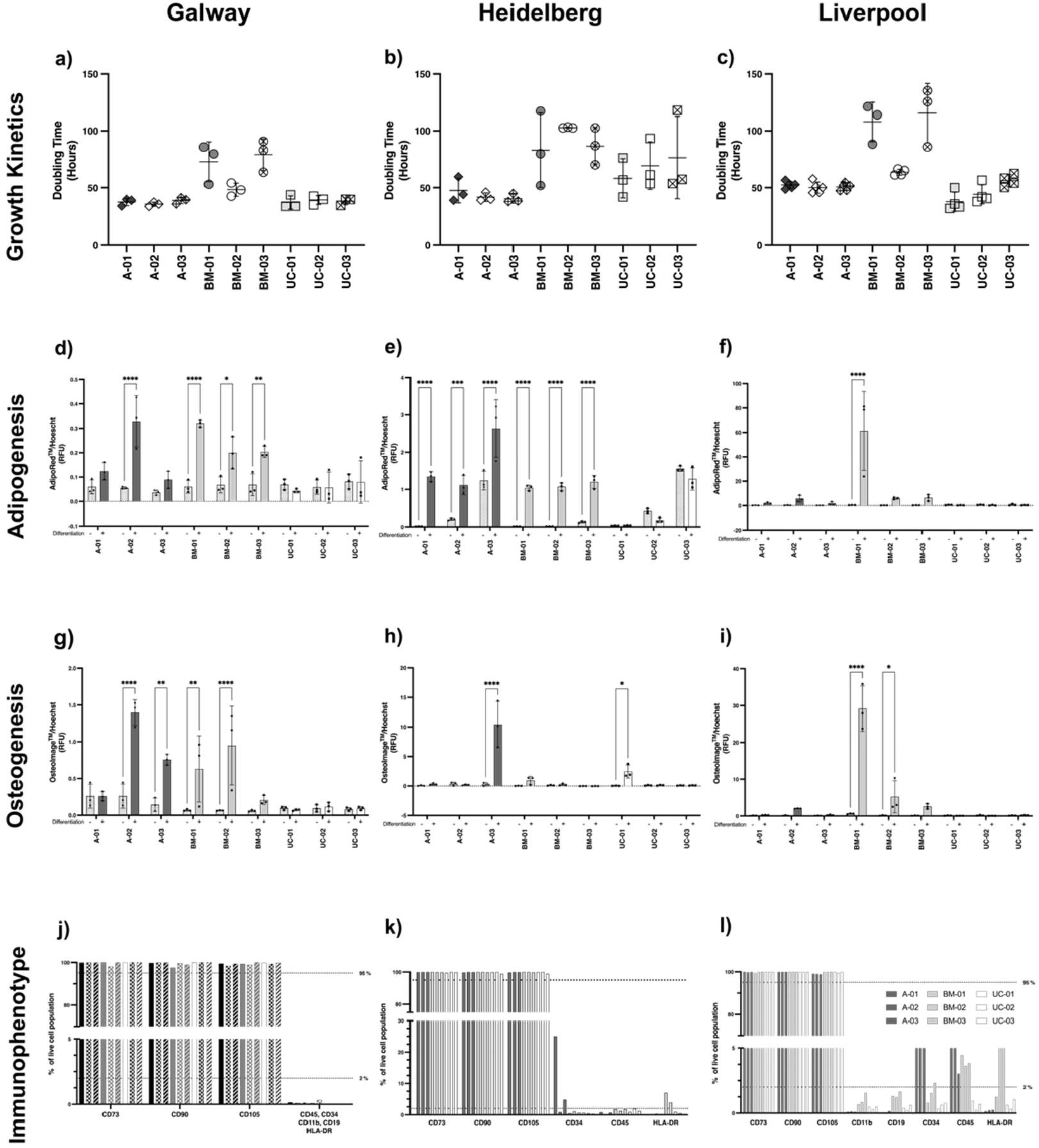
Biological comparison: donor-by-donor breakdown of doubling times, immunophenotype, differentiation results and phase contrast images of the differentiation. Figures (a-c) show the individual doubling times per each donor in all sites. The three dots within a single donor represent the doubling times from three consecutive passages. Across laboratories, A- and UC-showed stable proliferation rates when looking at individual donors. Greater differences were seen in BM-in terms of donor-to-donor variability, although each donor behaved similarly regardless of manufacturing site. In terms of committing to mesodermal lineages, high variability of induction was seen across laboratories. Broadly, A- and BM-donors were able to undergo adipogenesis in two sites, apart from one particular donor that showed induction in all laboratories (d-f). Negligible levels of adipogenic differentiation were seen in UC-MSC cultures. Similarly, A- and BM-MSCs were able to undergo osteogenic differentiation in two out of three sites (g-i), albeit not all donors and at remarkable different rates; exclusively one UC-MSCs in one site showed moderate levels of osteogenesis. Assessment of surface antigen expression confirmed >95% levels of CD73, CD90 and CD105 in all donors across sites (j-l). However, two preparations of A-MSC showed higher than 2% levels of CD34 in two and CD45 in one site. Importantly, these were the same donors. Data displayed as mean ± SD, N=3, n=3. One-Way ANOVA with Tukey’s multiple comparison corrections, * = p < 0.05, ** = p < 0.001, ** = p < 0.0001, **** = p < 0.00001.

**Supplementary Figure 5.**
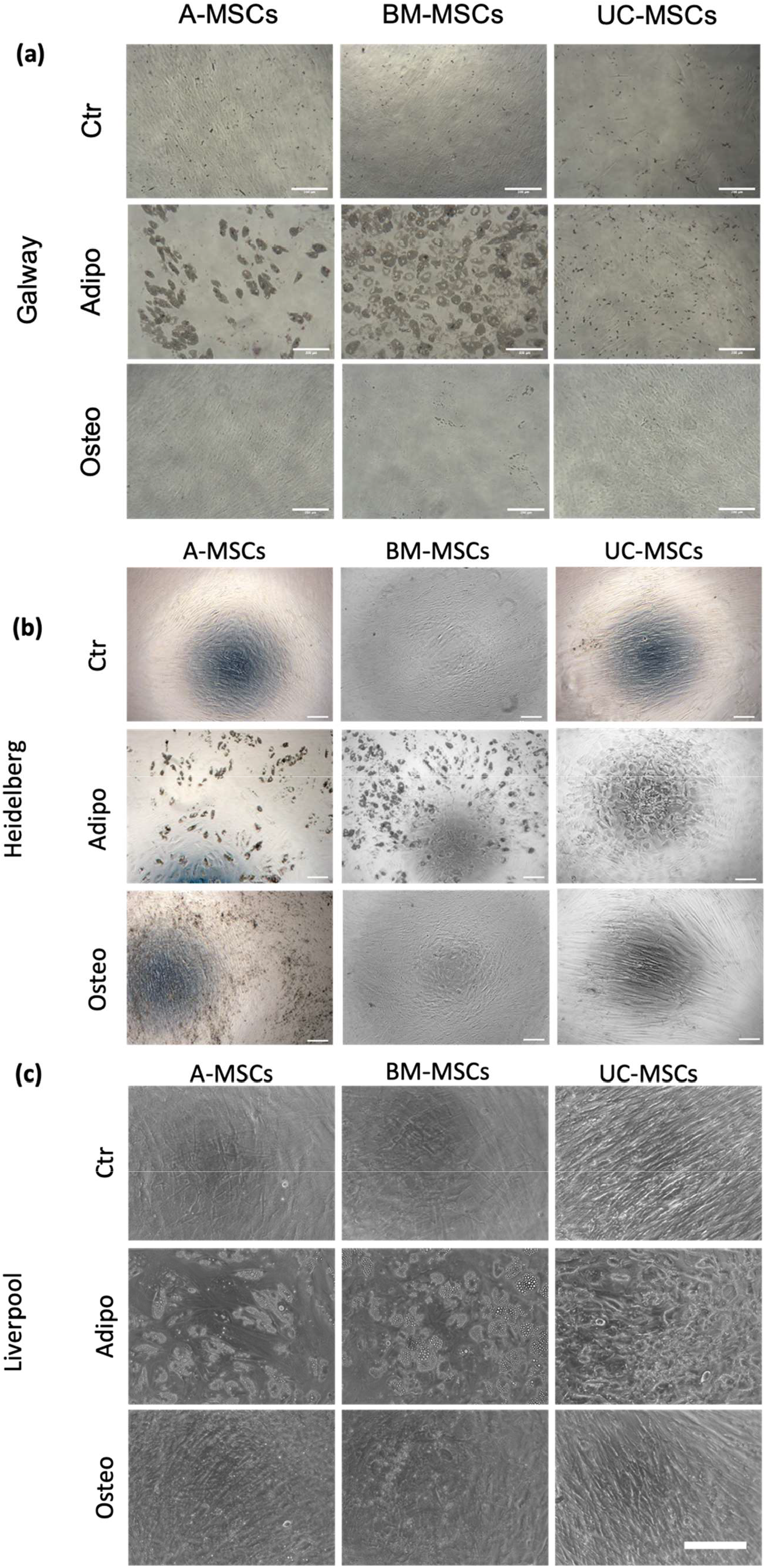
Representative phase contrast images of MSC at the end of the adipogenic (adipo) and osteogenic (osteo) differentiation procedure in comparison with undifferentiated cultures (ctr) in each site. Pictures taken at 100X; scale bar 200 µm.

**Supplementary Figure 6.**
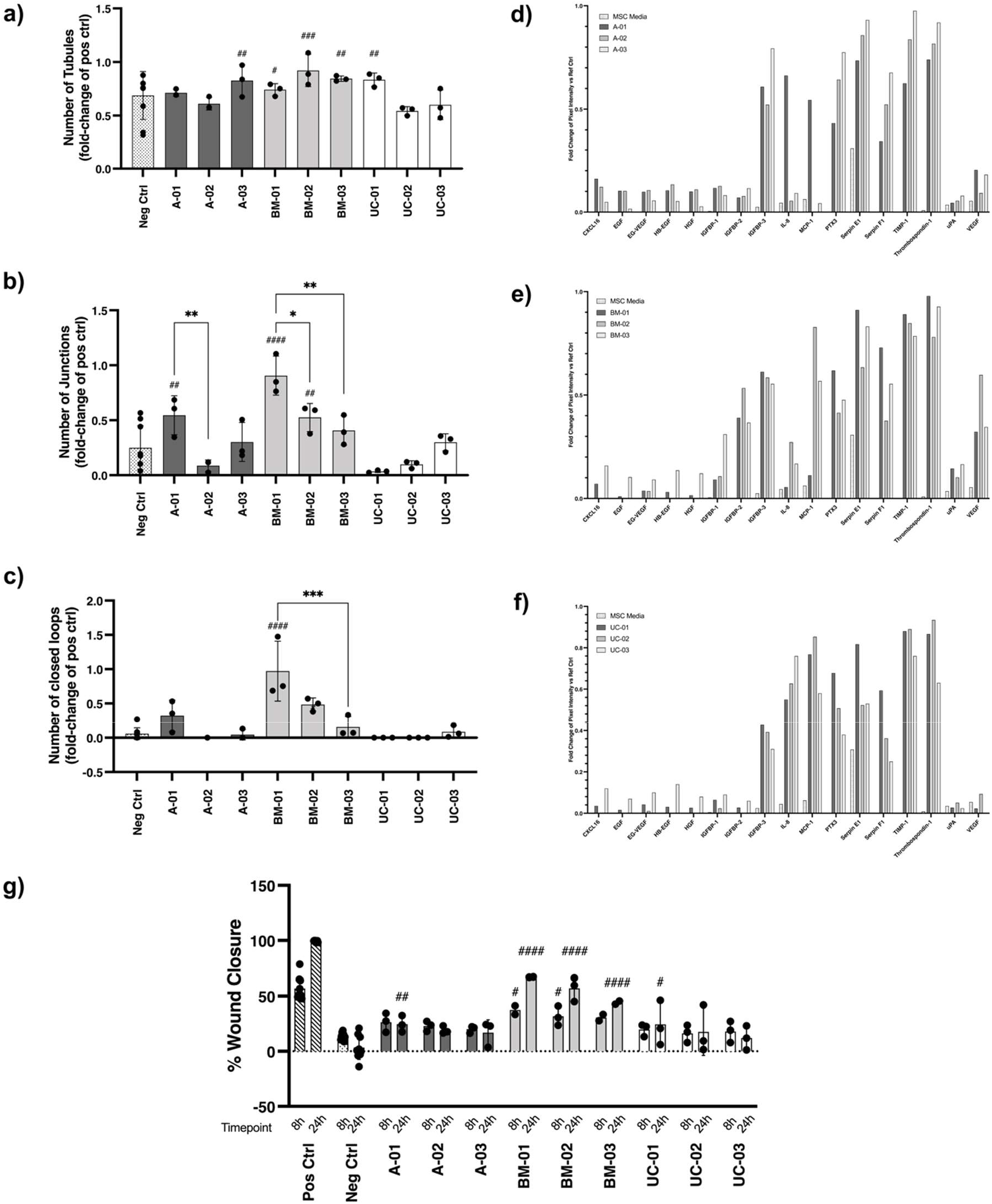
Angiogenic and wound healing properties of MSCs listed by donor. (a-c) Number of tubules (a), junctions (b) and closed loops (c) generated by each donor. Data expressed as a fold-change of the positive control; mean *±* SD, n = 3. (d-f) Differential angiogenic proteomic profile detected for A-(d), BM-(e) and UC-(f) MSCs. Data expressed as fold change of the internal reference spots. (g) Wound closure induced by each donor per cell source at 8 and 24 hours. Data displayed as mean *±* SD, n = 3. Two-Way ANOVA with Tukey’s multiple comparison corrections, * = p < 0.05, ** = p < 0.001, ** = p < 0.0001, **** = p < 0.00001. # Significance relative to negative control.

**Supplementary Figure 7.**
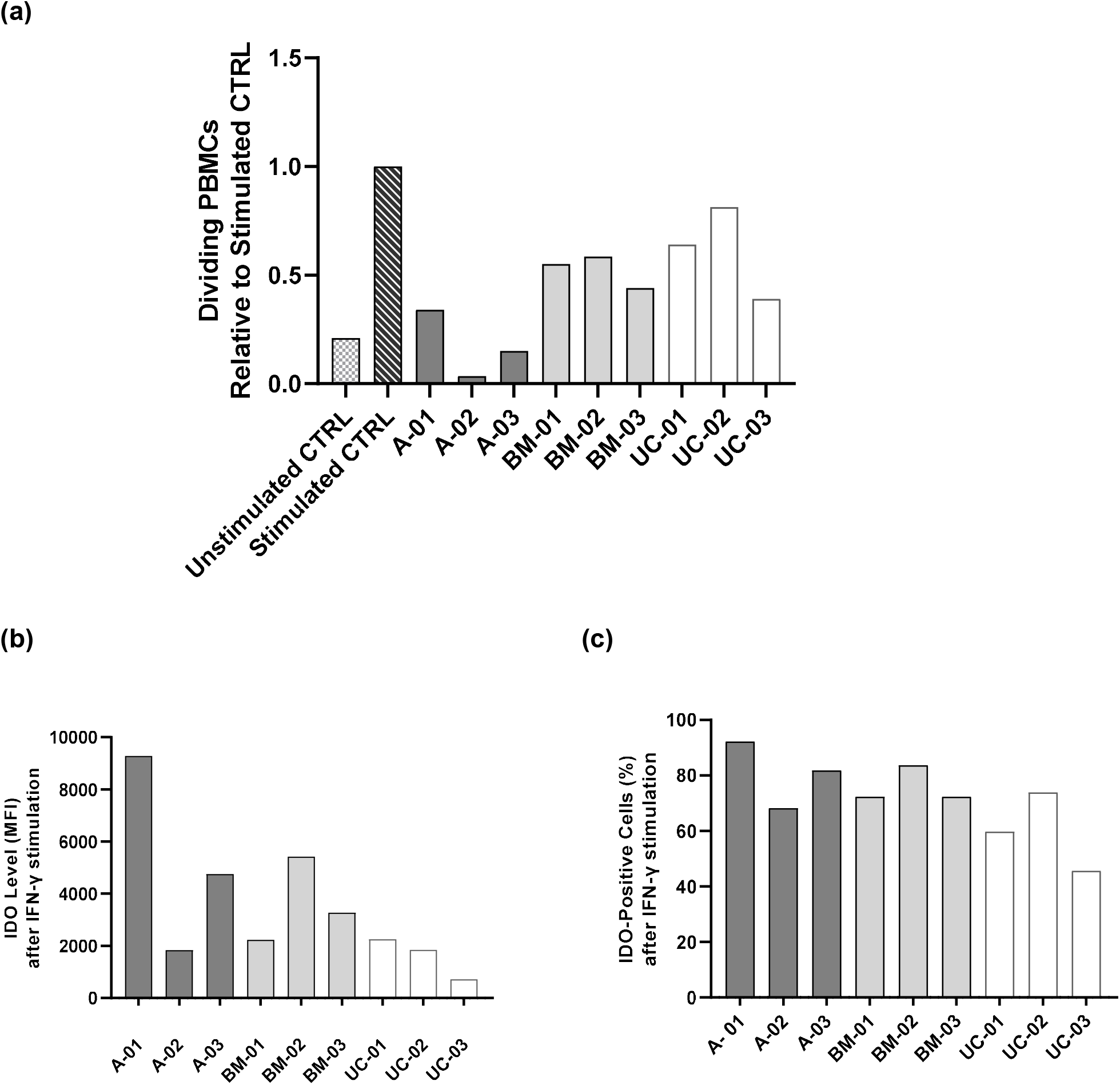
Donor-by-donor breakdown of MSC immunomodulatory capacity (a) Individual values of PBMC proliferation co-cultured with MSCs in the presence of PHA, where each bar represents the relative value in relation to PHA-stimulated PBMCs cultured alone. (b) MFI of IDO intracellular staining and (c) percentage of IDO-positive cells after 24h of IFN-γ stimulation,listed per donor.

**Supplementary Figure 8.**
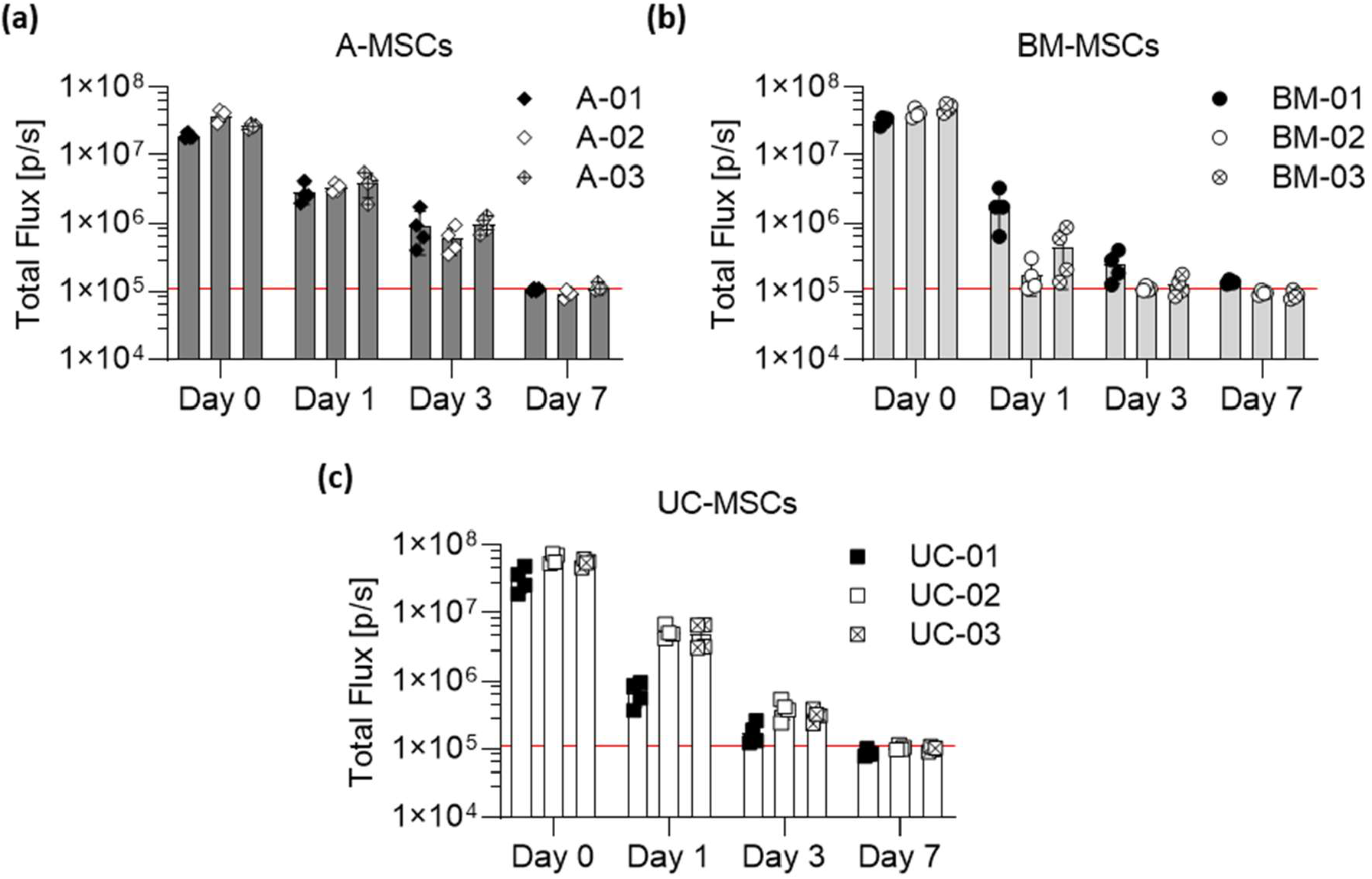
Donor-by-donor breakdown of the signal obtained from the *in vivo* imaging of MSCs in healthy C57BL/6 albino mice. (a-c) Light output (flux) as a function of time (days) coming from A-(a), BM-(b), and UC-(c) MSCs. Data displayed as mean ± SD from N = 4 for each donor. The red line (1.1 × 10^5^ p/s) is the background BLI signal emitted by naïve animals (n = 4) that did not receive any cells.

